# Compact type II-D Cas9 nucleases for efficient and specific genome editing

**DOI:** 10.64898/2026.08.17.745286

**Authors:** Qiaochu Wang, Ahmed Saleh, Gundra Sivakrishna Rao, Ahmed M. Kazlak, Rashid Aman, Magdy M. Mahfouz

## Abstract

Compact CRISPR nucleases are attractive for therapeutic genome editing because their small coding sequences facilitate delivery by adeno-associated virus. Type II-D Cas9 (Cas9d) enzymes constitute the most compact Cas9 subtype, yet only a few orthologs have demonstrated mammalian genome-editing activity, leaving it unclear whether this activity is general or exceptional. Here, we mined the IMG/M metagenomic database and identified five previously uncharacterized MG102-like Cas9d orthologs (∼950 amino acids) that share the hallmark genomic, sequence, and structural features of type II-D Cas9. Two of them, Cas9d-1 and Cas9d-4, recognized a 5’-NRC-3’ protospacer-adjacent motif and edited endogenous human loci with efficiencies up to 20.1%, exceeding *Streptococcus pyogenes* Cas9 at one site, while producing deletion-biased outcomes and no detectable off-target activity. Notably, both orthologs edited more efficiently than the sole previously validated member of this lineage, MG102-2, when assayed side by side under identical conditions. These findings establish compact MG102-like Cas9d orthologs as robust and specific genome editors and provide promising, single-AAV– compatible scaffolds for *in vivo* therapeutic genome editing.

## Introduction

Programmable genome editing with CRISPR–Cas9 has transformed biomedical research and is rapidly advancing toward clinical therapeutics,^1–5^ a transition crystallized by the recent approval of the first CRISPR-based medicine.^6^ Most of this progress rests on a single enzyme, the type II-A Cas9 from *Streptococcus pyogenes* (SpCas9), which pairs high activity in human cells with broad targeting through its short 5’-NGG-3’ protospacer-adjacent motif (PAM).^7^ Yet the features that made SpCas9 the field’s workhorse also constrain its therapeutic deployment: its 1,368-amino-acid coding sequence, together with a guide RNA and regulatory elements, approaches or exceeds the ∼4.7-kb packaging limit of adeno-associated virus (AAV), the leading *in vivo* delivery vector,^8, 9^ frequently necessitating dual-vector strategies that reduce co-transduction efficiency and complicate manufacturing.^10–13^ Identifying naturally compact nucleases that retain efficient, specific editing is therefore a central goal for next-generation genome editors.^14^

This search has yielded a remarkable diversity of compact RNA-guided enzymes. Compact Cas9 orthologs such as SaCas9,^15^ CjCas9,^16^ and the high-fidelity Nme2Cas9^17^ enabled single-AAV delivery, while class 2 type V systems including Cas12a,^18^ CasX (Cas12e),^19^ and the hypercompact Cas12f1^20^ have broadened the repertoire of compact programmable nucleases. Many were uncovered by systematic mining of microbial genomes and metagenomes,^21^ while the recognition that the transposon-encoded proteins IscB and TnpB function as programmable nucleases revealed the evolutionary origins of class 2 CRISPR systems and expanded the repertoire of naturally compact editing scaffolds.^22, 23^ Together with base and prime editing,^24, 25^ these advances have made compactness, specificity, and broad targetability defining design criteria for therapeutic editors.^26^

Type II-D Cas9 (Cas9d) sits at the extreme compact end of this spectrum. It is the smallest naturally occurring Cas9 subtype identified to date and is thought to be an evolutionary intermediate between canonical Cas9 and the transposon-encoded IscB family,^22, 27^ a uniquely informative position for both engineering and mechanistic study. Relative to type II-A, II-B, and II-C enzymes, Cas9d proteins carry a drastically shortened REC domain while retaining the conserved RuvC and HNH domains that catalyze RNA-guided DNA cleavage.^27–32^ They are further distinguished by enriched RRXRR and zinc-binding ribbon motifs and an arginine- and lysine-rich, methionine-depleted composition,^28^ a larger tracrRNA scaffold, and a recently described staggered-cut cleavage mechanism.^28, 29^ This combination of minimal size and ancestral character makes Cas9d both an appealing chassis for compact editors and a window into Cas9 evolution.

Despite this promise, the subtype remains under-exploited, with functional validation in human cells reaching only a handful of representatives. Two major groups have been identified: the longer MG102 lineage (∼950 aa) and the shorter MG34 lineage (∼750 aa).^28^ Subsequent characterization has focused largely on the MG34-like lineage, including MG34-1,^30, 31^ the related NsCas9d^29^ and NbaCas9^27, 32^ orthologs, and the reconstructed ancestor ancCas9d.^31^ By contrast, the longer MG102-like branch has been represented in mammalian cells by a single enzyme, MG102-2, leaving a fundamental question unresolved: is genome-editing competence an intrinsic property of this lineage, or a fortunate exception confined to one ortholog?^28^ Resolving it is a prerequisite for treating the MG102-like branch as a deployable source of compact editors.

Here, we address this question through discovery rather than engineering. Using a CRISPR-array–centered mining strategy applied to the IMG/M metagenome database, we recovered five previously uncharacterized MG102-like Cas9d orthologs and characterized them biochemically and in human cells. Two orthologs, Cas9d-1 and Cas9d-4, are bona fide RNA-guided nucleases that recognize a compact 5’-NRC-3’ PAM and edit multiple endogenous human loci reproducibly, with deletion-biased outcomes, no detectable off-target activity at predicted sites, and, at one locus, editing that exceeds SpCas9. By raising the number of mammalian-active MG102-like enzymes from one to three, these results establish genome editing as a general property of the longer Cas9d lineage and define it as a promising, AAV-compatible source of compact editors.

## Materials and Methods

### Identification of compact type II-D Cas9 candidates

A CRISPR-array–centered mining strategy was applied to metagenomic assemblies from the IMG/M database. CRISPR arrays were identified with CRISPRCasFinder 4.3.2,^33^ flanking regions (±15 kb) were extracted, and protein-coding genes were predicted with Prodigal^34^ and annotated against a custom hidden Markov model database of CRISPR-associated proteins. Non-redundant candidates (90% identity clustering; 400–1,000 aa) were retained after filtering for genomic context and locus completeness with CRISPRCasTyper.^35^ Candidates were aligned with representative type II-A–II-D Cas9 proteins (MAFFT v7.490;^36^ trimAl v1.5.1^37^) and placed in maximum-likelihood phylogenies with IQ-TREE v3.1.1^38^ (ModelFinder;^39^ 1,000 ultrafast-bootstrap and SH-aLRT replicates), visualized in iTOL.^40^ Catalytic-residue conservation and Cas9d sequence hallmarks were assessed by alignment; the RRXRR domain (PF14239) was detected with InterProScan,^41^ structures were predicted with AlphaFold3^42^ and compared to MG102-2 in PyMOL v3.1.8 (Schrödinger, LLC), and tracrRNAs were identified by a homology-guided strategy with RNA secondary-structure prediction (RNAfold).^43^ Full parameters are provided in Supplementary Methods (Supplementary Data S1).

### Plasmid construction and sgRNA design

Cas9d open reading frames were human-codon-optimized, synthesized (Twist Bioscience; Supplementary Table S1), and cloned into the pX330 backbone (Addgene #52970) under a CMV promoter; the SpCas9 plasmid (Addgene #87108) was obtained directly. sgRNAs were designed by linking each predicted crRNA repeat to its cognate tracrRNA through a 5’-GAAA-3’ tetraloop. For *in vitro* assays, templates bearing a 20-nt spacer matching the PAM-library protospacer^44^ were transcribed with the HiScribe T7 kit (NEB) and purified (Zymo Research). The sequences of sgRNA expression templates and oligonucleotides are listed in Supplementary Tables S2 and S3; detailed cloning and transcription procedures are in Supplementary Methods.

### Mammalian-lysate purification and in vitro cleavage assays

Cas9 expression plasmids were transfected into HEK293T cells (FuGENE 4K), and clarified soluble lysates were prepared 48 h later; FLAG-tagged protein expression was confirmed by Western blot. RNP complexes (lysate plus *in vitro*-transcribed sgRNA) were assembled and incubated with circular PAM-library substrate or XmnI-linearized target plasmid carrying a 5’-NRC-3’ PAM, and cleavage products were resolved on agarose gels. Buffer compositions, reagent amounts, and incubation conditions are detailed in Supplementary Methods; *in vitro* target sequences are in Supplementary Table S4.

### In vitro PAM screen

PAM preferences were determined with an N -randomized PAM library (Addgene #160132).^44^ Cleaved fragments were selectively enriched as described,^45^ ^46^ sequenced on an Illumina MiSeq, and the top 10% of enriched 8-bp PAMs (normalized to a no-guide control) were visualized as sequence logos with Logomaker.^47^ Library-preparation details and primers are in Supplementary Methods and Supplementary Table S5.

### Mammalian genome-editing assays

Guide RNAs targeting *CLTA*, *HBB*, *AIFM*, *Casp-3*, and *EMX1* (Supplementary Table S6) were cloned into U6-driven, ortholog-specific sgRNA vectors (Supplementary Table S7). HEK293T cells were co-transfected with Cas9 and U6-sgRNA cassettes (FuGENE 4K), genomic DNA was harvested at 72 h, and editing was assessed by T7 endonuclease I (T7E1) assay and amplicon deep sequencing. Cell-culture, transfection, and primer details are in Supplementary Methods and Supplementary Tables S8.

### Amplicon deep sequencing and statistical analysis

Target loci (Supplementary Table S9) were amplified in three successive PCR rounds using primers listed in Supplementary Table S10, sequenced on an Illumina MiSeq, merged with FLASH, and analyzed with CRISPResso2 v2.3.2.^48^ Editing below 0.1% or detected in untransfected controls was treated as background. Data are presented as mean ± s.e.m. of three biological replicates; group comparisons used the Kruskal–Wallis test with Dunn’s correction or two-way ANOVA with Bonferroni’s correction (GraphPad Prism v11), as indicated in the figure legends.

### Off-target prediction and analysis

Off-target sites were predicted with Cas-OFFinder v3.0.0^49^ against GRCh38/hg38 (up to four protospacer mismatches; 5’-NRC-3’ PAM; Supplementary Table S11), amplified using primers from Supplementary Table S12 and analyzed by deep sequencing as for on-target sites. Editing was scored as detectable only above both the untransfected control (mean + 3 s.d.) and a 0.1% threshold.

### Data and code availability

All next-generation sequencing data generated in this study have been deposited at the NCBI Sequence Read Archive (SRA) under BioProject accession PRJNA1484012 (SRA: PRJNA1484012) and are publicly available as of the date of publication. Plasmids constructed in this study are available from the corresponding author on reasonable request. All other data supporting the findings of this study are available within the article and its supplementary information.

## Results

### Mining of IMG/M metagenomes recovers five compact type II-D Cas9 candidates

To identify putative type II-D Cas9 nucleases, we surveyed protein neighborhoods of high-confidence CRISPR loci. CRISPR arrays called from IMG/M assemblies (>4 × 10 contigs) with CRISPRCasFinder^33^ yielded 823,181 arrays; protein-coding genes within ±15 kb were predicted with Prodigal,^34^ producing more than 9,000 Cas9-associated proteins. After clustering at 90% identity and filtering by length (400–1,000 aa), contig completeness, and locus architecture, 203 non-redundant candidates remained.

Maximum-likelihood phylogeny against representative type II-A–II-D Cas9 proteins placed a subset firmly within the type II-D clade, alongside the lineages described by Goltsman et al.^28^ After removing candidates with IscB hallmarks (most notably the conserved PLMP motif^22, 32^), we screened the remaining proteins for sequence and structural characteristics frequently associated with Cas9d enzymes, including the six conserved RuvC/HNH catalytic residues, enriched RRXRR and zinc-binding ribbon motifs, and arginine/lysine enrichment with methionine depletion.^28, 30, 31^ Five candidates met every criterion and clustered tightly with MG102-2, sharing ∼50% amino-acid identity with it (Fig. 1A); ^28^ these were designated Cas9d-1 through Cas9d-5 (Supplementary Data S1).

**FIG. 1.**
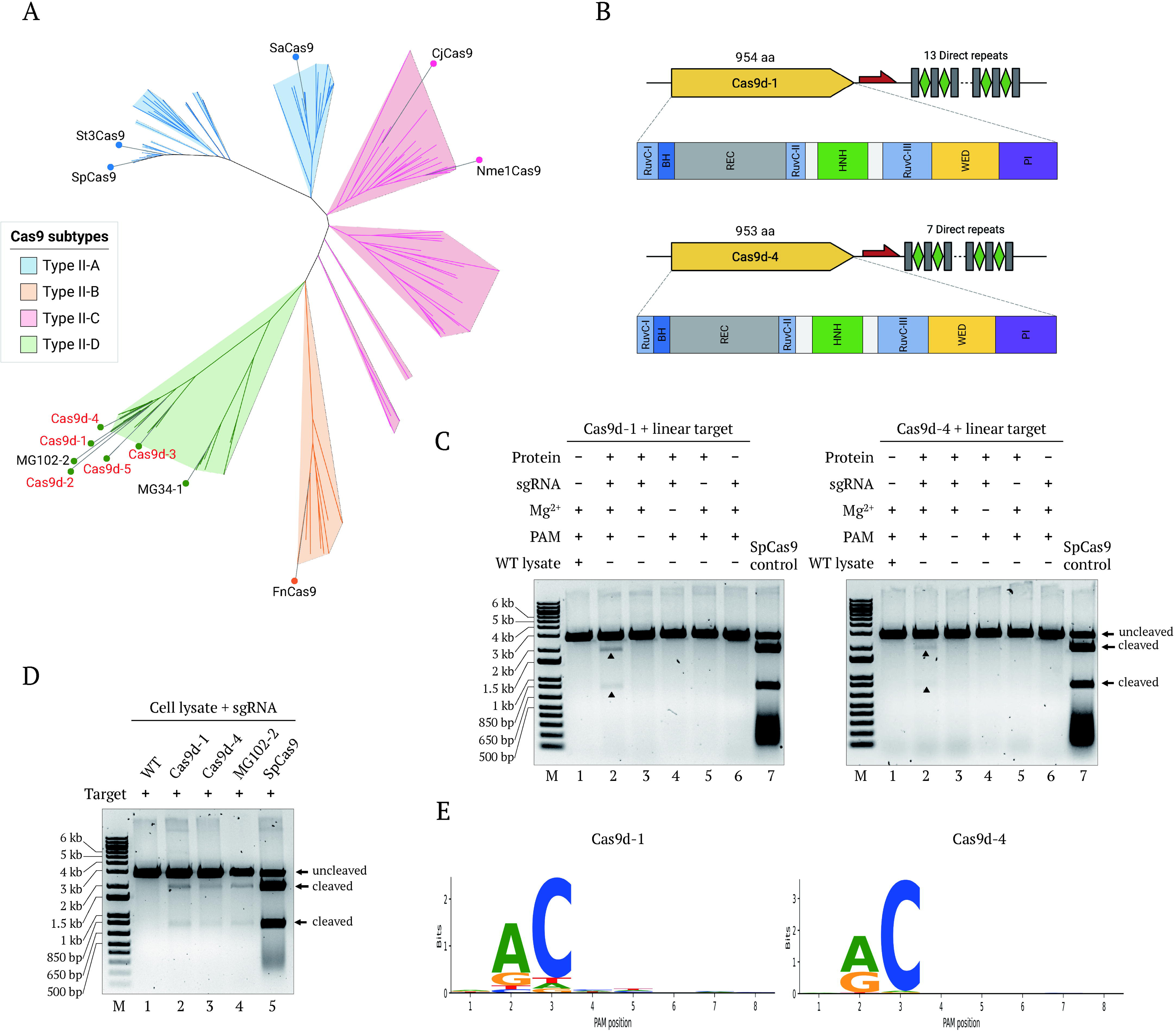
Identification and biochemical characterization of active type II-D Cas9 orthologs. **(A)** Maximum-likelihood phylogenetic tree of representative type II-A, II-B, II-C, and II-D Cas9 proteins. Newly identified Cas9d candidates (Cas9d-1 to Cas9d-5; red) cluster within the type II-D clade together with the previously characterized Cas9d proteins MG34-1 and MG102-2. **(B)** Genomic organization and predicted domain architecture of the two active orthologs, Cas9d-1 (954 aa) and Cas9d-4 (953 aa). Both loci contain a CRISPR array adjacent to the Cas9 open reading frame but lack the adaptation genes *cas1* and *cas2*. Predicted protein domains, including RuvC, bridge helix (BH), REC, HNH, WED, and PAM-interacting (PI) domains, are shown below each locus. Domain boundaries were inferred from sequence and structural alignments. **(C)** *In vitro* cleavage assays of linearized target plasmids by Cas9d-1 (left) and Cas9d-4 (right). Purified proteins expressed in HEK293 cells were assembled with *in vitro*-transcribed sgRNAs and incubated with target DNA under the indicated conditions. DNA cleavage required protein, sgRNA, Mg², and a compatible PAM sequence. SpCas9 was included as a positive control. **(D)** *In vitro* cleavage activity in a cell-lysate-based assay. HEK293 cell lysates expressing Cas9d-1, Cas9d-4, MG102-2, or SpCas9 were incubated with sgRNA and target plasmid DNA. WT lysate served as a negative control. **(E)** PAM preferences of Cas9d-1 and Cas9d-4 determined by *in vitro* screening of an N -randomized PAM library followed by deep sequencing. Sequence logos indicate a shared 5′-NRC-3′ PAM preference.

All five candidates were ∼950 aa and encoded immediately adjacent to CRISPR arrays but, like many type II-D loci, lacked the adaptation genes *cas1* and *cas2* (Fig. 1B, Supplementary Fig. S1A). Putative tracrRNAs were identified near each Cas9 locus, with predicted secondary structures resembling known Cas9d tracrRNAs (Supplementary Fig. S1B). AlphaFold3 models (pTM > 0.7) aligned closely with MG102-2 (backbone RMSD < 3 Å), adopting the characteristic bilobed Cas9 architecture (REC, RuvC, HNH, WED, and PI domains) with a markedly reduced REC domain (<300 aa) (Supplementary Fig. S2). Together, these genomic, sequence, and structural signatures classify Cas9d-1 through Cas9d-5 as bona fide type II-D Cas9 enzymes.

### Mammalian-lysate purification identifies two active Cas9d orthologs

To prioritize orthologs likely to function in human cells, we assessed *in vitro* DNA cleavage using proteins produced in a mammalian context, reasoning that bacterial activity does not always translate to human cells. Cas9d-1 through Cas9d-5 were human-codon-optimized, expressed from CMV cassettes in HEK293 cells, and purified from soluble lysates. Reconstituted with *in vitro*-transcribed sgRNA and incubated with circular or linearized target plasmid, only Cas9d-1 and Cas9d-4 showed sgRNA-dependent cleavage of both substrates, whereas Cas9d-2, Cas9d-3, and Cas9d-5 were inactive despite confirmed expression (Fig. 1C, Supplementary Fig. S3). Both active enzymes were less efficient than SpCas9 but comparable to MG102-2 (Fig. 1D), and were advanced for detailed characterization.

### Cas9d-1 and Cas9d-4 share a compact 5’-NRC-3’ PAM

To define PAM requirements *in vitro*, lysate-purified proteins were assembled with 20-nt sgRNAs and challenged with an N -randomized PAM library.^44^ Sequence-logo analysis of enriched, deep-sequenced products revealed a shared 5’-NRC-3’ PAM (R = A/G): minimal preference at position 1, strong purine selection at position 2, and a cytosine at position 3 (Fig. 1E). This closely matches MG102-2, consistent with the high conservation of their PAM-interacting domains, and the relaxed NRC motif is expected to occur frequently across the genome, affording broad target accessibility.

### Cas9d-1 and Cas9d-4 edit endogenous human loci with 20-nt spacers

To test editing in human cells, we selected ten endogenous sites across five genes (*CLTA*, *HBB*, *AIFM*, *Casp-3*, and *EMX1*; two each), each flanked by a 5’-NRC-3’ PAM. Co-expressing Cas9d with U6-driven sgRNAs in HEK293 cells, we first compared spacer lengths at three loci (*CLTA* target-1, *CLTA* target-2, and *HBB* target-1). Since the 20-nt spacers consistently outperformed 24-nt spacers that gave near-background editing (Fig. 2), all subsequent experiments used 20-nt spacers.

**FIG. 2.**
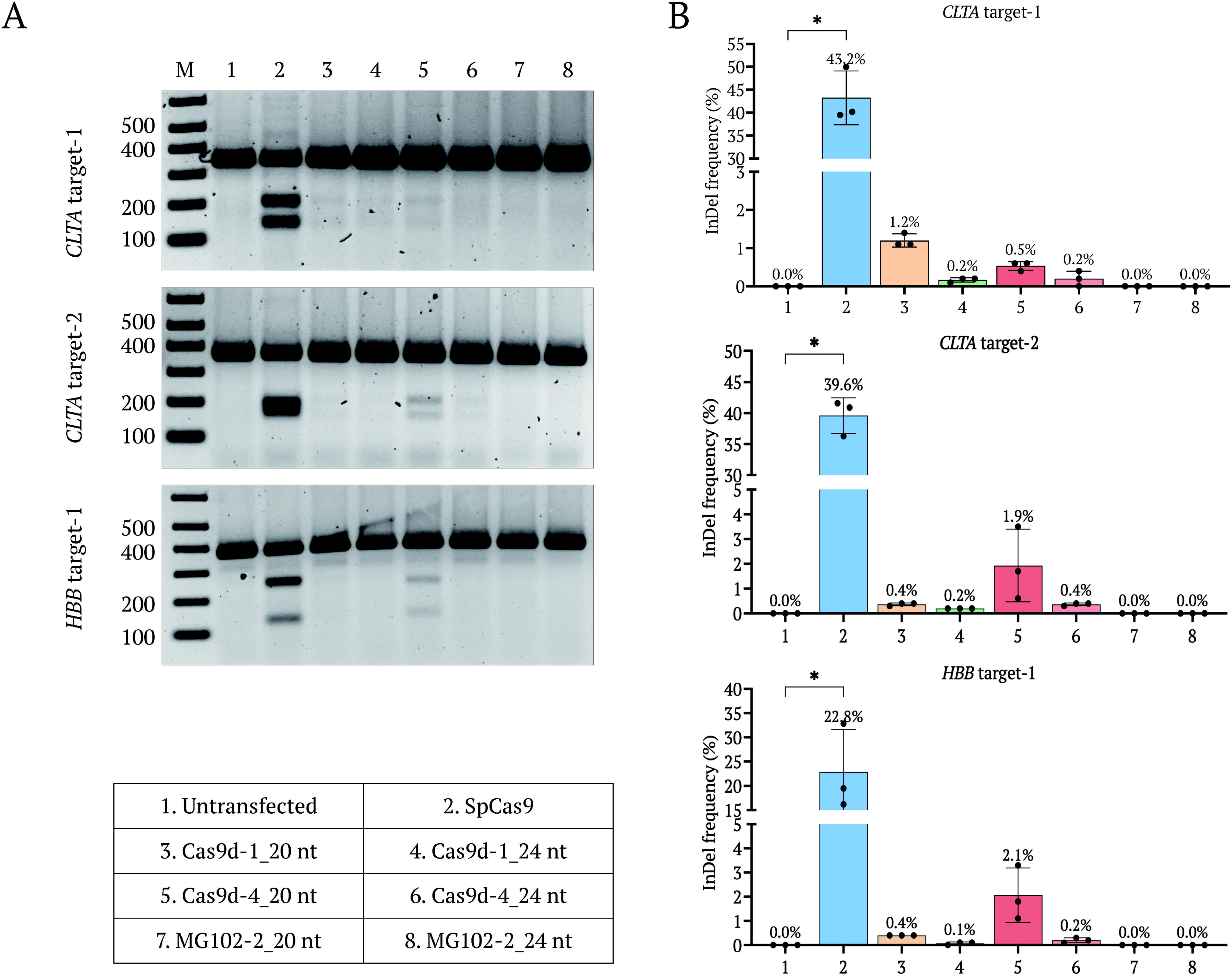
Comparison of spacer lengths for Cas9d-mediated genome editing at representative endogenous loci. **(A)** Representative T7 endonuclease I (T7E1) assays at *CLTA* target-1, *CLTA* target-2, and *HBB* target-1. Lane assignments: (1) untransfected control, (2) SpCas9, (3) Cas9d-1 with a 20-nt spacer, (4) Cas9d-1 with a 24-nt spacer, (5) Cas9d-4 with a 20-nt spacer, (6) Cas9d-4 with a 24-nt spacer, (7) MG102-2 with a 20-nt spacer, and (8) MG102-2 with a 24-nt spacer. M, DNA size marker. **(B)** Indel frequencies at the corresponding target sites determined by amplicon deep sequencing. Bars indicate mean editing frequencies and error bars represent s.e.m. from three biological replicates; individual replicate values are shown. Statistical significance was assessed using the Kruskal–Wallis test with Dunn’s multiple-comparison correction (*p < 0.05, **p < 0.01, ***p < 0.001, ****p < 0.0001).

With this architecture, we profiled the remaining sites alongside SpCas9 and MG102-2. Both Cas9d-1 and Cas9d-4 produced detectable editing across multiple loci by T7E1 assay and amplicon sequencing (Supplementary Fig. S4), with locus-dependent but reproducible efficiencies. Cas9d-1 was most active at *Casp-3* target-1 (20.1% indels) and *AIFM* target-2 (10.9%), and Cas9d-4 at *AIFM* target-1 (9.8%) and *AIFM* target-2 (8.4%). Across the ten loci, Cas9d-1 and Cas9d-4 edited seven and eight sites, respectively, at ∼1–20% (Fig. 3). Under identical conditions, the benchmark MG102-2 edited only a single site (*Casp-3* target-2; 0.6%), underscoring the substantially greater mammalian activity of both new orthologs. At *Casp-3* target-1, Cas9d-1 even exceeded SpCas9 (20.1% versus 10.8%), showing that naturally occurring Cas9d enzymes can outperform established editors at individual loci.

**FIG. 3.**
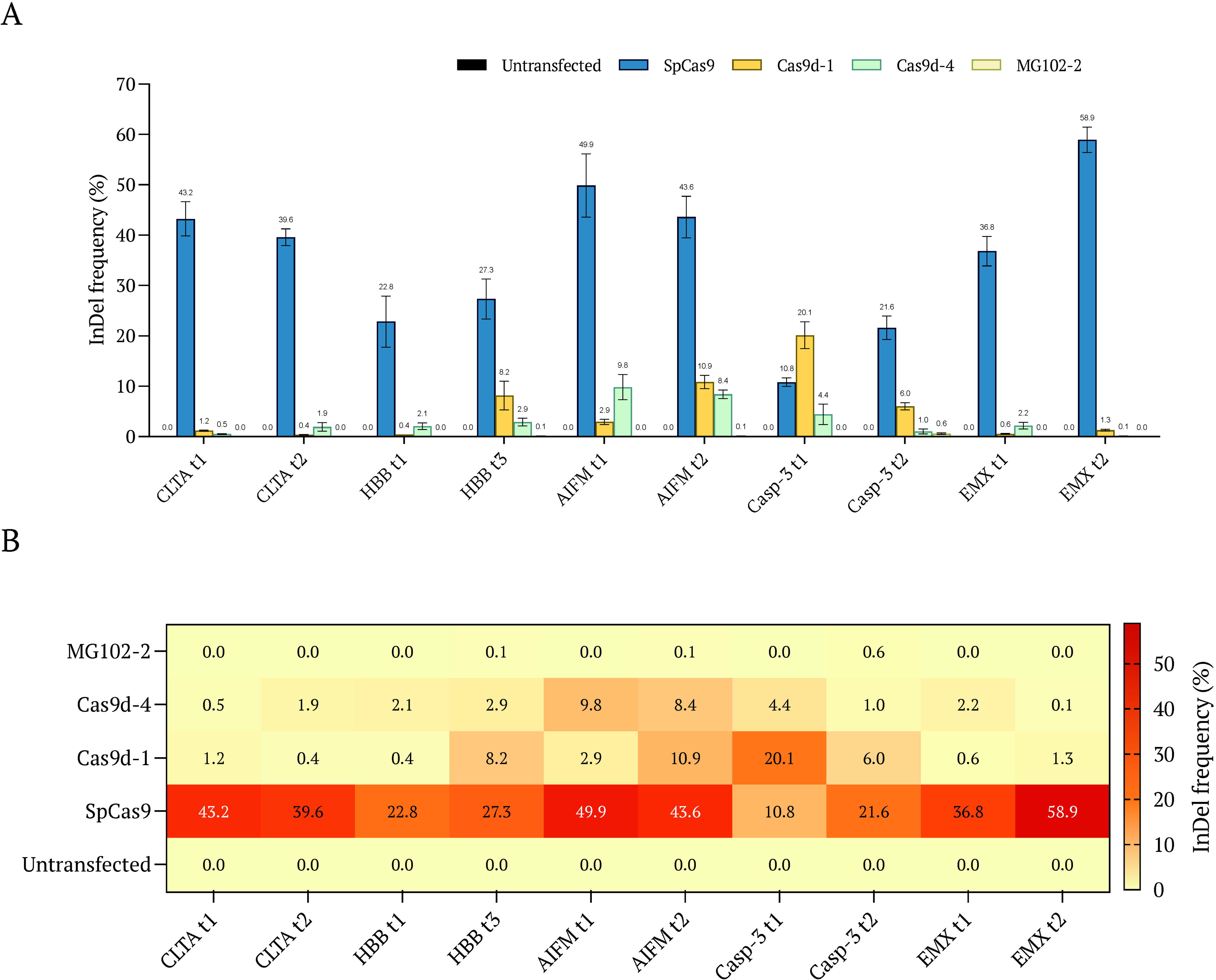
Genome-editing activities of Cas9d-1 and Cas9d-4 at endogenous human target sites using 20-nt sgRNAs. **(A)** Indel frequencies (%) at ten endogenous target sites determined by amplicon deep sequencing following transfection of HEK293 cells with SpCas9, Cas9d-1, Cas9d-4, or MG102-2 together with their corresponding sgRNAs. Editing frequencies were calculated after subtraction of background editing detected in untransfected controls. Bars represent mean ± s.e.m. from three biological replicates. **(B)** Heatmap summarizing mean indel frequencies for each nuclease across the ten endogenous target sites. Values within each cell indicate the mean editing frequency (%) obtained from three biological replicates.

### Cas9d editing yields deletion-biased mutation spectra and localized editing windows

To dissect editing outcomes, we examined four high-efficiency sites (*HBB* target-3, *AIFM* target-1, *AIFM* target-2, and *Casp-3* target-1). Both Cas9d-1 and Cas9d-4 generated predominantly deletions, with insertions a minor fraction, whereas SpCas9 produced a more balanced distribution (Fig. 4A). Pooled analysis across all ten sites confirmed markedly lower insertion-to-deletion ratios for Cas9d than SpCas9 (Supplementary Fig. S5), establishing deletion bias as a signature of Cas9d editing.

**FIG. 4.**
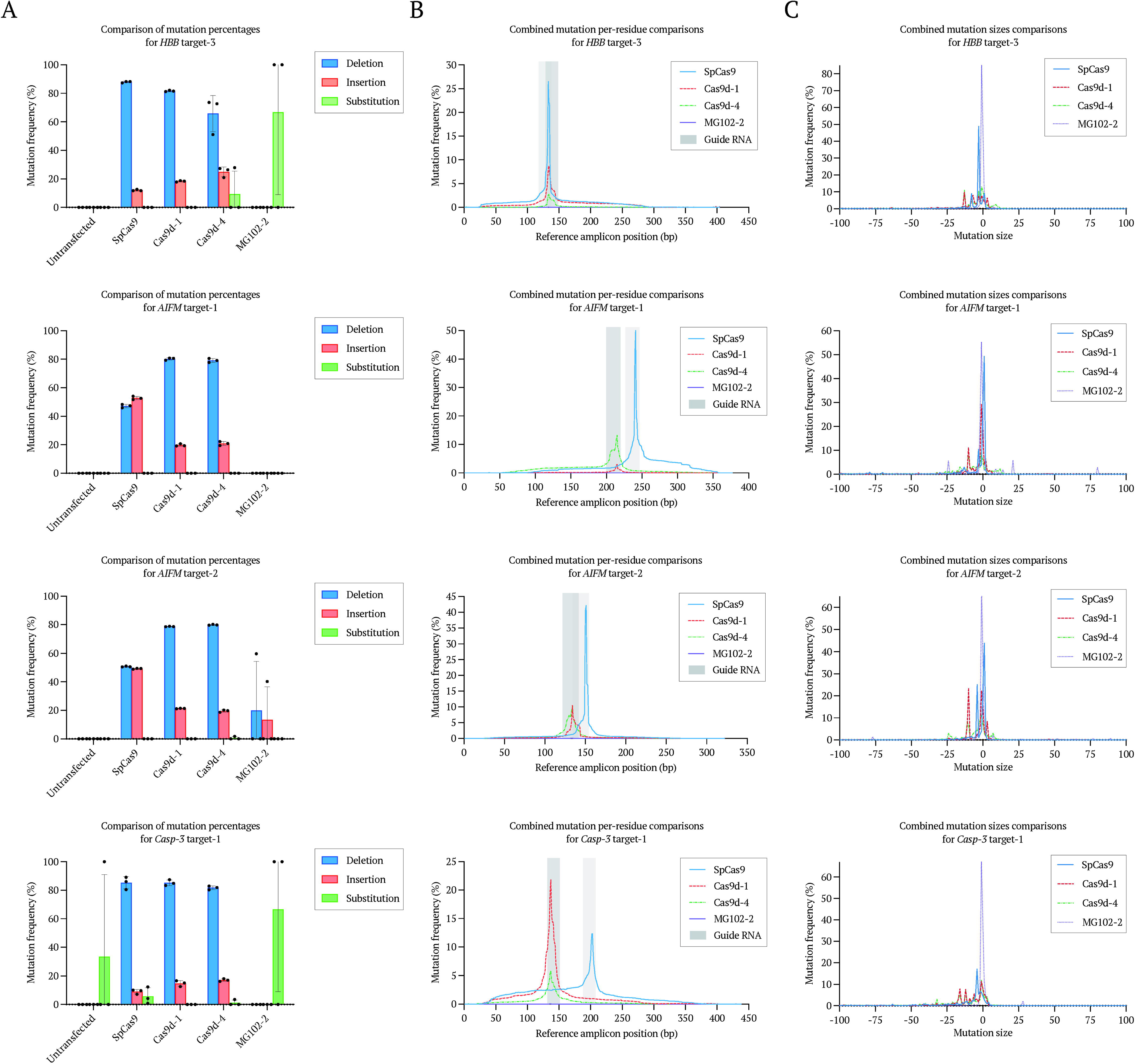
Mutation-outcome profiling of Cas9d-1 and Cas9d-4 at representative endogenous target sites. Composite mutation analysis at *HBB* target-3, *AIFM* target-1, *AIFM* target-2, and *Casp-3* target-1 across five conditions: untransfected, SpCas9, Cas9d-1, Cas9d-4, and MG102-2. **(A)** Relative proportions of deletions, insertions, and substitutions determined by amplicon deep sequencing. Mutation frequencies were calculated after subtraction of background events detected in untransfected controls. Values represent mean ± s.e.m. of three biological replicates, with individual replicate values shown. **(B)** Per-residue mutation frequencies across target amplicons, calculated from pooled insertion, deletion, and substitution events. Gray boxes indicate guide RNA target regions. Data represent the mean of three biological replicates. **(C)** Indel-size distributions at the four representative target sites. Positive values indicate insertions and negative values indicate deletions. Frequencies were calculated directly from amplicon sequencing reads without background subtraction and represent the mean of three biological replicates.

Per-residue profiling showed that Cas9d-1 and Cas9d-4 edits concentrated around the predicted cleavage site (Fig. 4B), remaining more spatially confined than SpCas9 edits across all four loci. Indel-size analysis confirmed a strong bias toward small deletions, mostly <20 bp (Fig. 4C). Notably, the two orthologs produced near-identical spectra at every locus despite their sequence divergence, suggesting a conserved cleavage-and-repair signature across the lineage.

### Specificity profiling reveals no detectable off-target activity at predicted sites

To assess specificity, we predicted off-target sites for the most active Cas9d-1 and Cas9d-4 guides with Cas-OFFinder,^49^ scanning GRCh38/hg38 for 5’-NRC-3’ sites with up to four protospacer mismatches. Eight sites for Cas9d-1 and nine for Cas9d-4 were analyzed by amplicon sequencing alongside on-target sites. At every predicted off-target, including sites differing by only two mismatches, indel frequencies were indistinguishable from untransfected controls (Fig. 5), indicating stringent on-target discrimination. Comprehensive genome-wide profiling by unbiased approaches (GUIDE-seq,^50^ CIRCLE-seq,^51^ and DISCOVER-seq^52^) will be valuable for determining whether this high specificity extends across the entire genome.

**FIG. 5.**
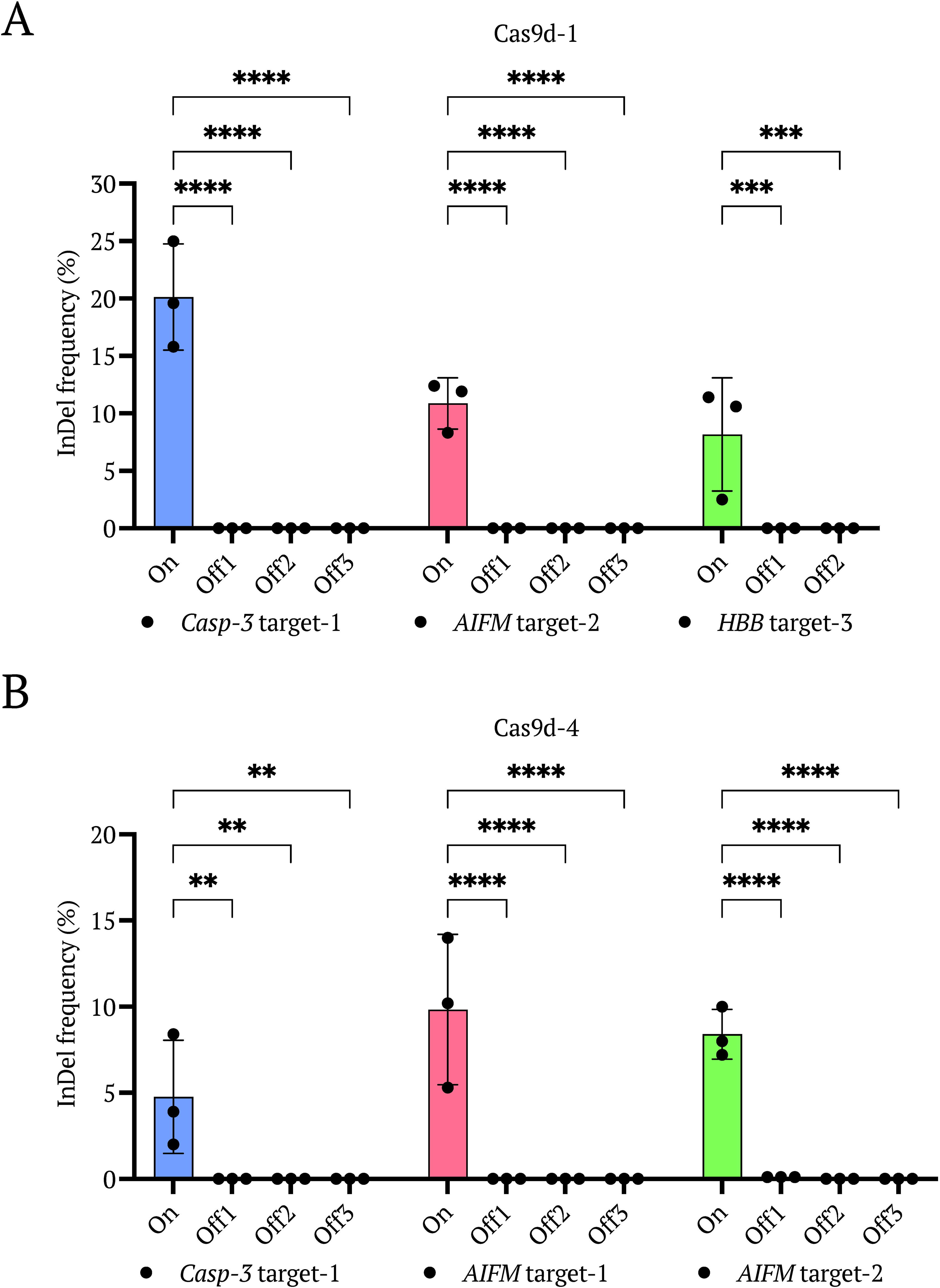
Off-target analysis of Cas9d-1 and Cas9d-4 at predicted genomic loci. Indel frequencies determined by amplicon deep sequencing at three representative on-target sites and their corresponding predicted off-target loci for Cas9d-1 **(A)** and Cas9d-4 **(B)**. Bars represent mean indel frequencies and error bars indicate s.d. from three biological replicates; individual replicate values are shown. Statistical significance was assessed using two-way ANOVA followed by Bonferroni’s multiple-comparison test. ns, not significant; *p < 0.05, **p < 0.01, ***p < 0.001, ****p < 0.0001.

## Discussion

By mining metagenomic data and characterizing five previously unreported MG102-like orthologs, we resolve a foundational question about the most compact Cas9 subtype: mammalian genome-editing activity is a general property of the longer type II-D lineage, not an idiosyncrasy of a single enzyme. Cas9d-1 and Cas9d-4 are close relatives of MG102-2, the only MG102-lineage ortholog previously validated in mammalian cells. They share the same lineage, ∼950-aa size, 5′-NRC-3′ PAM, tracrRNA architecture, and predicted fold; our contribution is therefore not the discovery of a structurally distinct scaffold, but rather the demonstration that mammalian genome-editing activity is a reproducible, conserved property of the MG102-like lineage rather than a unique feature of a single ortholog. Cas9d-1 reached 20.1% indels at *Casp-3* target-1 and exceeded SpCas9 there, demonstrating that an unengineered, naturally occurring ∼950-aa Cas9d can match or surpass the field’s benchmark editor at individual sites. This reframes the longer Cas9d branch as a deployable, AAV-compatible source of compact editors and positions natural-ortholog discovery, rather than engineering of known enzymes, as a fast route to expanding the toolbox. Beyond establishing activity, our parallel profiling of spacer-length preference, PAM recognition, editing across ten endogenous loci, mutation spectra, and predicted off-target sites constitutes, to our knowledge, the most comprehensive functional characterization of naturally occurring MG102-like Cas9d proteins reported to date.

Editing efficiencies varied substantially among targets, from background to ∼20%, consistent with the locus-dependent variation widely observed for CRISPR–Cas9 and commonly attributed to chromatin accessibility, local sequence context, and guide structure.^53, 54^ Importantly, activity at seven (Cas9d-1) and eight (Cas9d-4) of ten loci shows that editing is reproducible across genomic contexts rather than confined to a single favorable target. The benchmark MG102-2, highly active in its original report,^28^ edited only modestly here, likely reflecting differences in delivery (nucleofection versus FuGENE 4K), cargo (plasmid versus mRNA), cell line (HEK293 versus K562), and target selection, all of which can substantially influence editing.^55, 56^ Because all enzymes were benchmarked under identical conditions, the rankings are internally controlled, and matching the delivery format, particularly mRNA or RNP, should raise absolute efficiencies. A distinctive feature of these enzymes is their repair signature. Both orthologs produced strongly deletion-biased outcomes, with indels concentrated in short deletions (<20 bp) within a narrow window around the cleavage site. This predominance of deletions is consistent with the staggered DNA ends recently reported for type II-D Cas9,^28, 29^ and because DNA end structure influences double-strand-break repair-pathway choice, the staggered cut likely contributes to the observed bias.^57, 58^ The near-identical spectra of the two divergent orthologs suggest that deletion-biased repair is a shared property of the MG102-like lineage. A predictable bias toward small deletions is advantageous for loss-of-function editing and for therapeutic exon-skipping that disrupts splice donor or acceptor sites.^59^

The targeting range of Cas9d-1 and Cas9d-4 is set by a remarkably simple 5’-NRC-3’ PAM, expected to occur roughly once every eight base pairs per strand of random sequence and thus providing broad genomic access. Unlike the extended four-to six-base PAMs of many compact type II-C Cas9 orthologs, which restrict accessible targets while contributing to specificity, the short NRC motif imposes little constraint on targeting.^60^

Despite this permissive requirement, Cas9d-1 and Cas9d-4 showed no detectable editing at any computationally predicted off-target, including sites differing by only two mismatches. Although candidate-based analyses sample only part of the genome, the absence of editing at closely matched sites suggests strong discrimination; because neither an extended PAM nor a long guide accounts for it, this fidelity may be intrinsic to the Cas9d–guide RNA complex. Unbiased genome-wide approaches (GUIDE-seq, CIRCLE-seq, and DISCOVER-seq)^50–52^ will be needed to confirm whether this specificity is maintained genome-wide and preserved during engineering. Translationally, the appeal of the MG102-like lineage lies in coupling compact size with measurable, specific mammalian activity. At ∼950 aa, Cas9d-1 and Cas9d-4 are encoded by ∼2.85-kb open reading frames, roughly 30% smaller than SpCas9, leaving substantially more room within the ∼4.7-kb AAV cargo for a guide-RNA cassette, regulatory elements, or accessory effector domains, making them attractive scaffolds for compact base and prime editors, transcriptional regulators, and other tools.^61–63^

The near-identical behavior of two independently discovered orthologs indicates that the MG102-like lineage offers a robust, engineerable scaffold rather than an isolated example. As shown for other natural nucleases, activity and specificity can often be improved by guide-scaffold optimization, nuclear localization signal engineering, structure-guided design, and directed evolution.^64, 65^ High-resolution structures of Cas9d-1 or Cas9d-4 ribonucleoprotein complexes will help define PAM recognition and cleavage and guide engineering that preserves the compact architecture.

More broadly, this work shows that systematic metagenomic mining, coupled with sequence, structural, and functional characterization, is an effective strategy for expanding the natural diversity of genome-editing nucleases. A CRISPR-array–centered pipeline applied to a single database recovered two mammalian-active editors from an under-explored lineage, and the approach applies in principle to any RNA-guided nuclease class. Given the exponential growth of metagenomic sequence space, continued mining, together with the engineering and structural studies above, should yield additional compact editors with improved activity and broadened targeting.

## Conclusion

We report the discovery and biochemical and cellular characterization of five previously uncharacterized compact type II-D Cas9 nucleases, two of which, Cas9d-1 and Cas9d-4, mediate efficient, specific genome editing in human cells, with Cas9d-1 exceeding SpCas9 at an endogenous locus. By raising the number of mammalian-active MG102-like enzymes from one to three, these findings establish genome-editing activity as a general property of the longer Cas9d lineage rather than a single-enzyme trait. Together with their simple 5’-NRC-3’ PAM, deletion-biased outcomes, undetectable off-target activity at predicted sites, and AAV-compatible size, these enzymes provide a promising foundation for mechanistic study, protein engineering, and the development of compact editors for therapeutic genome editing in delivery-constrained settings.

## Supporting information

Cas9d_SI

Cas9d_Supplementary_Data_S1

## Authorship Confirmation Statement

M.M.M. conceived and supervised the project. Q.W., A.S., and G.S.R. designed and performed the experiments. A.M.K. performed the metagenomic mining and bioinformatic analyses. R.A. and A.S. assisted with cloning, mammalian transfections, and protein purification. Q.W., A.S., and M.M.M. drafted the manuscript with input from all authors. All co-authors have reviewed and approved the manuscript prior to submission. The authors confirm that this manuscript has been submitted solely to The CRISPR Journal and is not published, in press, or submitted elsewhere.

## Acknowledgments

This work was supported by King Abdullah University of Science and Technology (KAUST) baseline research funding (BAS/1/1035-01-01) to M.M.M. We thank the KAUST Bioscience Core Lab for next-generation sequencing services and the KAUST Supercomputing Laboratory for access to computational resources used for metagenomic mining and phylogenetic inference.

