## Supplementary material for "Compact type II-D Cas9 nucleases for efficient and specific genome editing": Cas9d_SI

**Supplementary Methods**

**Identification of compact type II-D Cas9 candidates**

A CRISPR-array-centered mining strategy was applied to metagenomic assemblies from the IMG/M database. CRISPR arrays were identified using CRISPRCasFinder 4.3.2,^1^ and genomic regions encompassing each array together with up to 15 kb of flanking sequence on both sides were extracted. Protein-coding genes within these regions were predicted using Prodigal^2^ and annotated using a custom hidden Markov model (HMM) database comprising CRISPR-associated proteins and accessory factors. To reduce redundancy, proteins were clustered at 90% sequence identity and the longest sequence from each cluster was retained as the representative. Candidate proteins were further filtered by length (400–1,000 aa) and genomic context, excluding proteins located within 200 bp of contig termini or loci exhibiting evidence of truncation or incomplete CRISPR-Cas architectures based on CRISPRCasTyper^3^ annotations.

The remaining candidates were analyzed together with representative type II-A, II-B, II-C, and II-D Cas9 proteins (Supplementary Data S1). Protein sequences were aligned using MAFFT v7.490^4^ (L-INS-i, 1,000 iterations) and trimmed with trimAl v1.5.1^5^ (-automated1). Maximum-likelihood phylogenetic trees were constructed using IQ-TREE v3.1.1^6^ with the optimal substitution model selected by ModelFinder.^7^ Branch support was estimated using 1,000 ultrafast bootstrap and 1,000 SH-aLRT replicates, and trees were visualized using iTOL v7.^8^

Candidates clustering within the type II-D Cas9 lineage were subjected to further sequence and structural analyses. Conservation of catalytic residues and characteristic Cas9d sequence features was evaluated manually by multiple-sequence alignment. The RRXRR domain (PF14239) was detected using the InterProScan online server.^9^ Protein structures were predicted using AlphaFold3^10^ and compared with the experimentally characterized Cas9d nuclease MG102-2^11^ using PyMOL v3.1.8 (Schrödinger, LLC). Putative tracrRNAs were identified using a homology-guided strategy integrating sequence similarity to published MG102 tracrRNAs, anti-repeat identification, and RNA secondary-structure prediction (RNAfold web server, Andronescu energy model (2007)^12^). Candidates satisfying phylogenetic, sequence, structural, and genomic criteria were selected for experimental characterization (Supplementary Data S1).

**Plasmid construction and sgRNA design**

Each Cas9d open reading frame was human-codon-optimized and synthesized (Twist Bioscience) as a cloning insert (Supplementary Table S1), and cloned into the pX330 backbone (Addgene #52970) under the CMV promoter, replacing the original FokI-Cas9 ORF; the SpCas9 plasmid (Addgene #87108) was obtained directly. Single-guide RNAs (sgRNAs) were designed by linking the predicted crRNA repeat to the cognate tracrRNA through a 5′-GAAA-3′ tetraloop, generating repeat duplexes of 15–20 bp. For the *in vitro* cleavage assays, sgRNA transcription templates containing a 20-nt spacer complementary to the fixed protospacer in the PAM library^13^ were cloned into a T7-driven sgRNA expression vector (Addgene #160136). Templates were amplified by PCR, transcribed *in vitro* using the HiScribe T7 High-Yield RNA Synthesis Kit (NEB, #E2040S) and purified with the RNA Clean & Concentrator-25 kit (Zymo Research) according to the manufacturers' instructions. The sgRNA templates and amplification primers are provided in Supplementary Tables S2, S3.

**Mammalian-lysate purification and *in vitro* cleavage assays**

HEK293T cells were maintained in DMEM (1×) GlutaMAX-I high-glucose medium without sodium pyruvate (Gibco, #61965-026) supplemented with 10% heat-inactivated fetal bovine serum (HI-FBS; Gibco, #16140-071) at 37 °C in a humidified incubator with 5% CO₂. Cells were seeded at 1.5 × 10⁶ cells per well in 6-well plates and transfected the following day with 2 µg of Cas9 expression plasmid using 13.4 µL FuGENE 4K Transfection Reagent (Promega, #E5911) in 72 µL Opti-MEM I Reduced Serum Medium (Gibco, #31985-062). Forty-eight hours after transfection, cells were harvested and lysed in buffer containing 50 mM Tris-HCl (pH 7.5), 0.1% NP-40 (Thermo Scientific, #85124), and 10% glycerol supplemented with cOmplete EDTA-free protease inhibitor cocktail (Roche, #11873580001). Lysates were clarified by centrifugation at 13,000 × g for 15 min at 4 °C, and the soluble fractions were aliquoted and stored at −80 °C. Protein expression was verified by Western blotting using a monoclonal ANTI-FLAG M2 antibody (Sigma-Aldrich, #F3165).

For *in vitro* cleavage assays, RNP complexes were assembled by incubating 20 µL of clarified lysate with 2 µg of *in vitro*-transcribed sgRNA and 1 µL RNaseOUT Recombinant Ribonuclease Inhibitor (Invitrogen, #10777019) at 37 °C for 15 min. Cleavage reactions were initiated by adding either 400 ng of circular plasmid substrate containing the cognate protospacer flanked by an N₈-randomized PAM library (Addgene #160132) or 300 ng of XmnI-linearized target plasmid containing a 5’-NRC-3’ PAM in 1× reaction buffer (10 mM Tris-HCl, pH 7.5, 100 mM NaCl, 10 mM MgCl₂, and 1 mM DTT). Reactions were incubated at 37 °C for 1 h, followed by treatment with RNase A (2 mg/mL; NEB, #T3018L) for 15 min at 37 °C and proteinase K (2 mg/mL; Thermo Scientific, #EO0491) for 15 min at 50 °C. Cleavage products were resolved by electrophoresis on 0.9% agarose gels. Sequence details for the *in vitro* targets are listed in Supplementary Tables S4.

**In vitro PAM screen**

PAM preferences were determined using an N₈-randomized PAM library (Addgene #160132)^13^ containing a fixed protospacer flanked by an 8-bp randomized PAM sequence. In vitro cleavage reactions were performed by incubating 50 µL of preassembled RNP complexes with 1 µg of PAM library plasmid in a total volume of 100 µL 1× reaction buffer at 37 °C for 1 h. Cleaved DNA fragments were selectively enriched for deep sequencing as previously described with minor modifications^14^ ^15^. Briefly, cleavage products were end-repaired with 1 U T4 DNA Polymerase (NEB, #M0203S), dA-tailed using 3 U TaKaRa Ex Taq DNA Polymerase (TaKaRa, #RR001A), and ligated to a 3′-dT adapter. Adapter-ligated fragments were enriched by two rounds of PCR, during which Illumina sequencing adapters and sample-specific indices were incorporated, and were subsequently sequenced on an Illumina MiSeq platform. Sequencing reads were filtered for a minimum Phred quality score of 20, aligned to the plasmid backbone, and the randomized 8-bp PAM sequences were extracted. The top 10% most abundant PAM sequences were retained, normalized to a no-guide control library, and visualized as sequence logos using Logomaker.^16^ Sequence details for the adapters and primers used for deep sequencing are listed in Table S5.

**Mammalian genome-editing assays**

Guide RNAs targeting endogenous sites within the *CLTA*, *HBB*, *AIFM*, *Casp-3* (*Caspase-3*), and *EMX1* genes were designed and synthesized as complementary oligonucleotides by Integrated DNA Technologies (Supplementary Table S6). Guide sequences were cloned into U6-driven sgRNA expression vectors containing the corresponding ortholog-specific scaffold, synthesized by Twist Bioscience (Supplementary Table S7), and U6-sgRNA cassettes were amplified by PCR for transfection. The primers used for the sequencing and amplification of sgRNA cassettes are listed in Supplementary Table S8. HEK293T cells were seeded at a density of 8 × 10⁴ cells per well in 0.1% gelatin-coated 24-well plates and transfected at approximately 70% confluency using FuGENE 4K Transfection Reagent (Promega, #E5912) with 250 ng Cas9 expression plasmid and 250 ng U6-sgRNA cassette per well. Cells were harvested 72 h after transfection, and genomic DNA was isolated using the Monarch Genomic DNA Purification Kit (NEB, #T3010L). Genome-editing activity was initially evaluated by T7 endonuclease I (T7E1) assays (NEB, #M0302L) using PCR amplicons spanning each target locus (Supplementary Table S9), followed by amplicon deep sequencing as described below.

**Amplicon deep sequencing and statistical analysis**

Target loci were amplified by three successive rounds of PCR to introduce locus-specific sequences, Illumina adapter overhangs, and sample indices, using primers listed in Supplementary Table S10. Libraries were pooled and sequenced on an Illumina MiSeq platform. Paired-end reads were merged using FLASH and analyzed with CRISPResso2 (v2.3.2).^17^ Editing frequencies detected in untransfected controls or below 0.1% were considered background and excluded from downstream analyses. Data are presented as mean ± s.e.m. from three biological replicates unless otherwise indicated. Statistical analyses were performed using GraphPad Prism v11.0.0. Comparisons among multiple groups were performed using the Kruskal–Wallis test followed by Dunn’s multiple-comparison test or two-way ANOVA followed by Bonferroni’s multiple-comparison test, as indicated in the corresponding figure legends. No statistical methods were used to predetermine sample size, and experiments were not randomized or performed in a blinded manner.

**Off-target prediction and analysis**

Potential off-target sites were identified using Cas-OFFinder (v3.0.0)^18^ against the human reference genome (GRCh38/hg38), allowing up to four mismatches within the protospacer and requiring a compatible 5’-NRC-3’ PAM (Supplementary Table S11). Selected off-target loci were amplified and analyzed by amplicon deep sequencing using the same workflow as for on-target sites, using primers listed in Supplementary Table S12. Editing events were considered detectable only when indel frequencies exceeded both the corresponding untransfected control (mean + 3 s.d.) and the empirical detection threshold of 0.1%.

**Supplementary Figures**

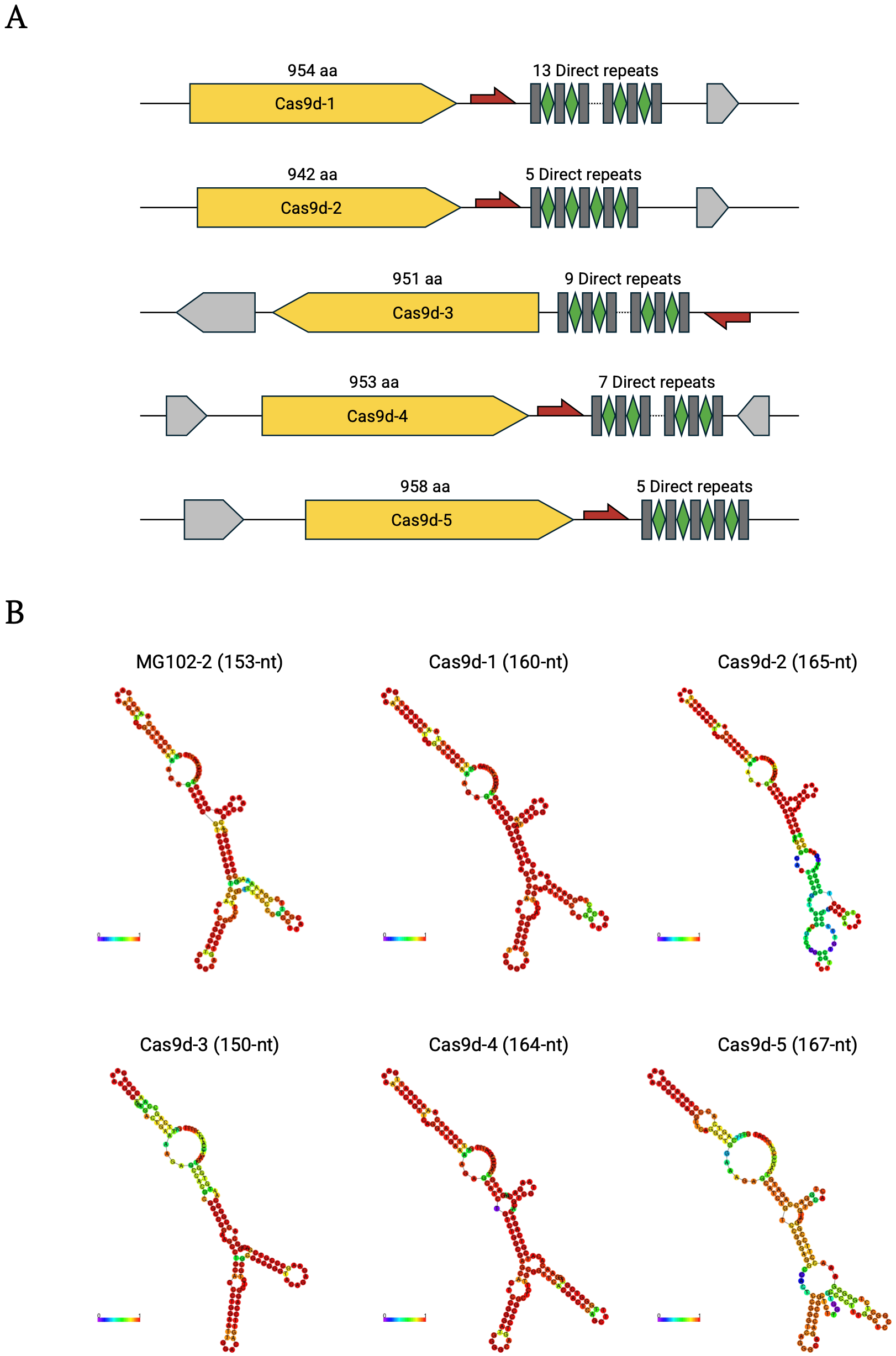

**FIG. S1. Genomic organization and predicted tracrRNA secondary structures of the five Cas9d candidates.**

**(A)** Genomic organization of the five Cas9d candidate loci. The Cas9 open reading frame (yellow), predicted tracrRNA (red), CRISPR direct repeats (gray rectangles) and spacers (green diamonds), and neighboring open reading frames (gray arrows) are shown. Protein lengths and the number of CRISPR direct repeats associated with each locus are indicated. **(B)** Predicted secondary structures of the corresponding sgRNA scaffolds generated using the RNAfold web server^12^ with the Andronescu energy model (2007)^19^. The predicted tracrRNAs range from 150 to 167 nt in length and display conserved stem-loop architectures characteristic of type II-D CRISPR-Cas systems. Scaffold of MG102-2 was used as control.

**FIG. S2. AlphaFold3 structural prediction of the active Cas9d orthologs Cas9d-1, Cas9d-4 and the control Cas9d MG102-2.**

**(A)** Predicted three-dimensional structures of Cas9d-1, Cas9d-4 and MG102-2 generated by AlphaFold3. The canonical Cas9 domains, including the recognition (REC), bridge helix (BH), RuvC, HNH, WED, and PAM-interacting (PI) domains, are indicated. **(B)** Predicted structures colored according to per-residue confidence (pLDDT). Regions with lower confidence were primarily confined to flexible loop regions and inter-domain linkers. **(C)** Predicted aligned error (PAE) matrices for Cas9d-1, Cas9d-4 and MG102-2 generated by AlphaFold3. Darker colors indicate lower predicted alignment error and higher confidence in the relative positioning of residue pairs.

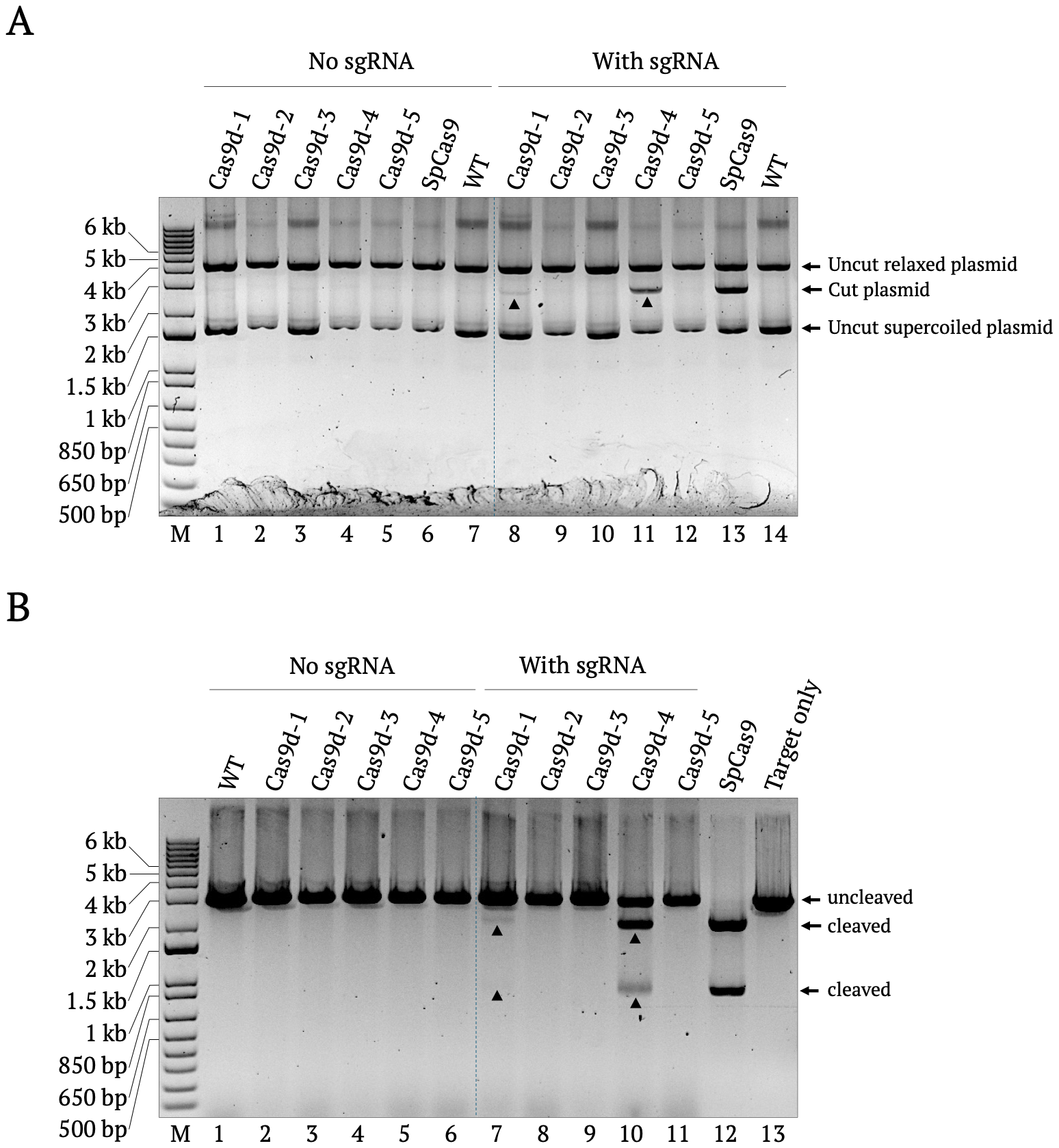

**FIG. S3. Initial *in vitro* screening of Cas9d candidates using mammalian cell lysate-derived proteins.**

**(A)** *In vitro* cleavage assay using a circular plasmid substrate containing an N₈-randomized PAM library. Cas9d-1 to Cas9d-5 and SpCas9 were transiently expressed in HEK293T cells, and clarified mammalian cell lysates were incubated with *in vitro*-transcribed sgRNA to assemble RNP complexes. Cleavage reactions were performed in the absence or presence of sgRNA. WT, lysate prepared from untransfected HEK293T cells. The positions of the supercoiled, relaxed, and cleaved plasmid forms are indicated. Cas9d-1 and Cas9d-4 exhibited detectable sgRNA-dependent cleavage activity, whereas Cas9d-2, Cas9d-3, and Cas9d-5 showed no detectable cleavage under the assay conditions. **(B)** *In vitro* cleavage assay using an XmnI-linearized plasmid containing a canonical 5’-NRC-3’ PAM. Reaction conditions were identical to those in (A). Cleaved and uncleaved DNA fragments are indicated. Consistent with the circular plasmid assay, Cas9d-1 and Cas9d-4 exhibited sgRNA-dependent cleavage activity, whereas Cas9d-2, Cas9d-3, and Cas9d-5 showed no detectable activity. SpCas9 served as a positive control and WT lysate and reactions lacking sgRNA served as negative controls.

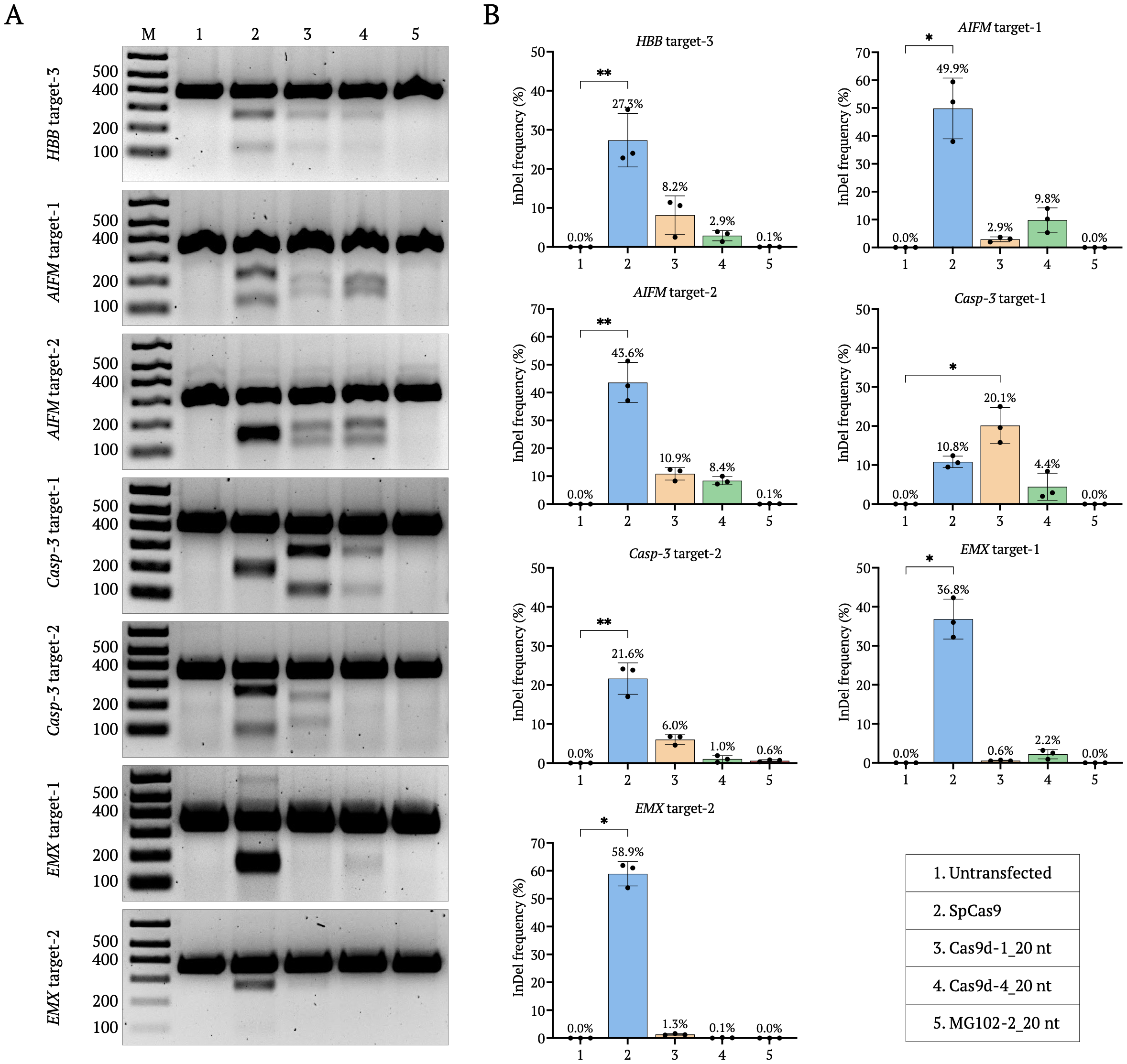

**FIG. S4. Genome editing by Cas9d-1 and Cas9d-4 at seven additional endogenous human target sites.**

**(A)** Representative T7 endonuclease I (T7E1) assays for ***HBB*** target-3, ***AIFM*** target-1, ***AIFM*** target-2, ***Casp-3*** target-1, ***Casp-3*** target-2, ***EMX1*** target-1, and ***EMX1*** target-2. Lane assignments are: M, DNA size marker; (1) untransfected control; (2) SpCas9; (3) Cas9d-1; (4) Cas9d-4; and (5) MG102-2. **(B)** Indel frequencies at the corresponding endogenous target sites determined by amplicon deep sequencing. Editing frequencies were calculated after subtraction of background editing detected in untransfected controls. Bars represent mean ± s.e.m. from three biological replicates, and individual replicate values are shown. Statistical significance was assessed using the Kruskal–Wallis test followed by Dunn's multiple-comparison test. P < 0.05, **P** < 0.01, **P < 0.001, ***P < 0.0001.

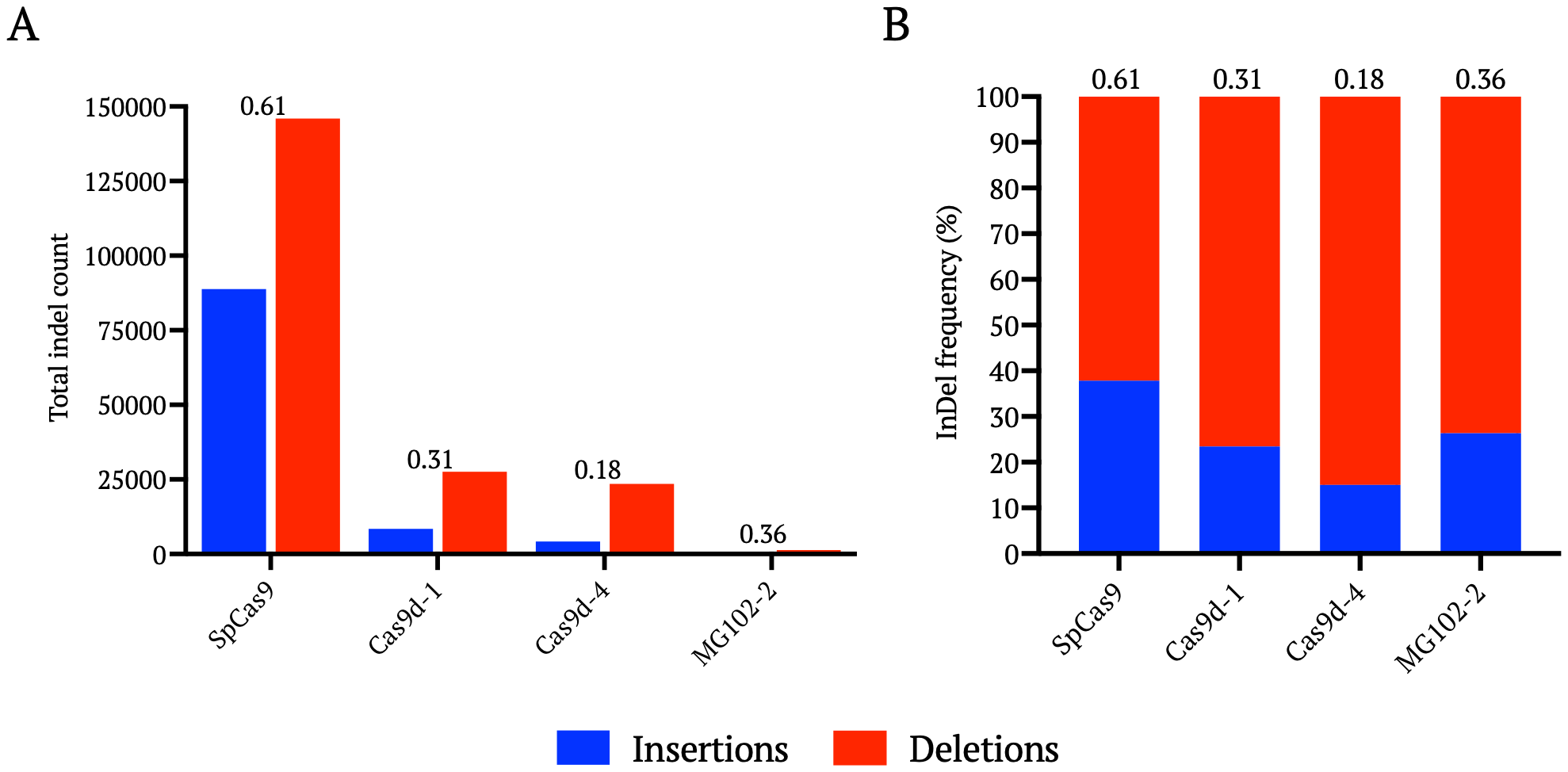

**FIG. S5. Genome-wide insertion-to-deletion profiles of Cas9d nucleases.**

Aggregate insertion and deletion events were calculated from amplicon deep-sequencing data across all analyzed endogenous target sites and biological replicates for SpCas9, Cas9d-1, Cas9d-4, and MG102-2. **(A)** Total insertion (blue) and deletion (red) counts for each nuclease. The insertion-to-deletion (Ins:Del) ratio is indicated above each group. **(B)** Relative proportions of insertions (blue) and deletions (red), normalized to 100% for each nuclease. The corresponding insertion-to-deletion (Ins:Del) ratios are shown above each bar. Both Cas9d-1 and Cas9d-4 exhibited lower insertion-to-deletion ratios than SpCas9, indicating a stronger preference for generating deletions following DNA cleavage.

**Supplementary Tables**

**Table S1. Human codon-optimized type II-D Cas9 coding sequences**

| **Name** | **Sequence (5’---3’)** |
| --- | --- |
| Cas9d-1 | AGT**GGATCC**ATGGAAAATATTCTGCAGACCGGCAGAACCTACACCCTGGGCATCGACTACGGCGCAAGTAATGTTGGTATTGCACTGGTGTGCAACGAGCCTACAGGCGAAAACCTGCCTCTGTTTGCAGCCACCCTGAGATTGGATGCACGGGATCTGAAAGAAAAAGTTGAGACACGTGCAGGTATTAGAAGCCTGCGTAGAACAAAGAAAACCAAGAAACATCGCCTGCATGACCTCGCAGATGCTCTGAAGAGCATCGGCCTGCCTGAACGTCAGGTTCAGAGCGTGATTCGTTTCAGCAAACGCAGAGGTTACAAGAGTCTGTTCGATGAAGATCTGCATGATGAAACCGAAGAGCAGCAGGACACCGATTTTGTGTATCGTTTTAGCCGTGAAGTGTTCTTCCAGTCTCTGAGCAGCGAACTGGAGAATATTATTGCCGACCCGGTTCTGAGACTGAATACGCTGGCCCAGTGTGAACGTATCCTGAACAAACAGGGCAACCCAGACCACGAAATCCGACGGATCAGAATCGACAATAGAGGCGCCTCCCGTTGCGCCTGGGCAGGATGCAATCGTGTGACACCTCGTTCCGATAACGCCATTCGTGACGTGCTGGCTCAGCAGCTGTATAACTACTTTCAGGCCCTGCTGAGGGACAACCCGAGCCAGAGCGCCAAAGTGGAAGATGTTATCCAAGAGCTGGTCGTAATCGCAAAACGGATTCGTGCAGCATCAGGCGATCACATCAAAGAGGAGAAGAAAGCGCTGCGTAAACGAGCACGTGGAGCGCTGCGGGGACTGAACAAGCTGCTGATGCCTGTTGCAACCGGCATCCCTGAAGATGAGCCTTGGGAATATGTTGAAAAGGGCATGATGAACACAATCGAGCGGCGTGCCGGACGTAATCGTTATTGTAGAGAGCACAGCGCAGAATACGTGCAGACCATCCTGGCCGGTAAAGCCGTTCCGTTTAAAAAGACCATCACCGACGCCGACATAATTAGCCGACGTGAACAGATGGCTTATAGCAAGCTGTGGCGCTACATCGAGGCCAGAATCCTGCCCCTGGCGCCTCAGGGCATTGATCGTATTGTGGTGGAACGTACCGCATTTGATCTGCTGGCAGGTAGCTGGAAGACCATCCAAGGTGCCACCGATAAATACAAAGAAGAGATGTATCAGCAAGGTCCGATGTTCGGTTTTAGCAGCAAGAGGGAAATGCTGAACGAGGAGTTCGGCGGTCTGTGTGCCTACTGCGGTAAACCGAGTGCTGATCTGATGGACCACGACCACATCATGCCGAGGGCTGATTTTTTCTTTGATGGCTATCTGAATATCTTACCGGCCTGTCCGACCTGTAATACCAACCTGAAAGGTAAACGTGCCCTGAGCGACCAAGCACTGACCATCGTGCCCGAGGCCTATGAAGCCTATAAGAAGTACCTGGCAACCAAGTTTCGCACCAGACCCATGCACTTCCTGCATACCGTGAAAAAAGGTATTTTGAATCTGATGAAAGATCCTGATAGAATCTGGGAGGCAGAAAAATATCTGAGCATTGTGGCCAGACAGTATGCACAGATCGTCCAGACACAGAGGAGCCCCCGTCCGTTCGCAAGGCTGCTATGCAGAAAAATCAAAGAAAGCCAGGGTTCCGAGCCCTCCATCGCATTCCGAAGCGGTCGGCATTCTGCTCTGTACAGACAGATTGCCTATCCGGAATTTGAAAAGTATGCTGATAAAGAACAAGGTAATGTGATCAACCACGGCCTGGACGCCATTCTGTTAGCATCTTCTCTGCCCGACCTGTATCCTGTGGAAAGCCTGAATATTCCTATCGCACGTCTGAAAGGCTGGACAGCCTCCGTGCGTAAGCGCGCACCGCGTCCAGGTAGCGATGGCATACCTGCAATTAGCGTTAAAGGTATGGTGGATGGGTTTGAAAAAATCCATGCCCCTGGATATATCGAGGTGGAAGTGCGTGCCATGGTTTGGAATAAAAAGAACAGCCGGACCCATAAACAGGACCCTTATGGTTGGTCCGAAAGCCATCACATGCCGACCAAACGCAAGGCCGCAGCAGATCTGTATGCCGATATGACTAAAGAAAACAGTCCTACCAGACTGAAGGCTATGGTGGAGCGGATATATCATCCAGCCCTGAAGCACTTCATTCTGGGCCGTATGGATACCGACAAACCGGGCCAGAGCGCCGCCGCCGCACTGAAAGAATGGCTGCAGGCAAGCATCCGTAATAGCCTGCCGGATAGCCAGTTTAGCAACCACCCTGGCGATCAGGCTCGTAAAGCGGATCTGGAAATGTTTGCCGAAGTTCCAGCCGCCCCGATTCCGGCAATTATTGGCGTGAAGATGATCGATACCGGCGTTCAGGGCAAAATCGACCTGGAACGTATGGATCATAAAACAGGACGGATTGTTCACAGGTATATGACAGACCCACCGAATAAAGCAGTTTATGTTGCCTACCCTAAGAAACGGGATGGTGGCTCCGATATGACCAAGCCTTGTCTGCTGTTTCTGCGTCAGAACGACGCCATCATTGTTGAAAGCCTGGCAGCATTTCGTCCGCTGCCCGAAGAACTGGCCAGAGGACGTGTCCTGGGTAGCACCCAGAGCAAGATTTTTGATGCGGCACTGCTGGAGAAATACCTAGAAGAATGTGGCTTCCACTGCTACATCCGGCTGACCAGCGGCTGCGTGATTAATTACGAGGATGGCAAAAGATGGTTCGTTAGGAACTTCGATCAGAGCAAAGACTTCAAAAAAGCGAGACTGCGGAACATCAAGGGTGTCCAGAAAACCCCTTTCAGCCGAAGAATCGAACCACTGGTGTACCTGGGTAAGTAATTAATTAAC**ACCGGT**ATC |
| Cas9d-2 | AGT**GGATCC**ATGATCACCCTGGGAATAGACTACGGCTCCAGCAATATTGGCATCGCCCTGGTTCGAAACACCGGCGAGGGCAACGAGCCTCTGTTTGCCGGCACCATCAAGATTGATGCGCGTCGTCTGCGTGATAAGGTGGAGACACGTGCAGGTATCCGGAGACTGCGACGCACGCGTAAAACCAAACGGCACCGTCTGCGGAACTTGCGCAACCGTTTTCTGTCTTTAGGTATTAAAGAGGAAGATGTTAATCTGATCGTGGGCTTCTGCAAGCGTAGAGGCTACAAGAGCCTGTTCGATGATGGTGAAACAGTGGAAAGCGAGAAAGGTGATAATCAGCTGACCTATCGTTTTACCAGGGAGGAATTCTTTCGTAACCTCACAGACGAGCTGAAGACCATCCTGCCTGACAGCAGCCAGTTTCAGGATGCACTGAACGCCTGTGAAAAAATTCTGAACAGACGGGGTGATCCGTTTCAAGAGATTCGTCTGATTAGGATCGATAACCGGGGCGCCAGCAGATGCGCCTGGGAAGGTTGTGATAAAGTGACCCCGAGAAGAGACAACGCACTGGGTGATCCGATTGCCCAGAGCCTGCTGAATGCAGTTCAAGAGAACCTGAAAGGGGAGCCTGCGAGAATGGGTACGATCGAGACCGACATTAGCGAACTCGATACACTGGGTAGACGCATTCGTGCAGCCTCTGGCGAAGGCGCTAAAGAAGAAAAGAAAGCCCTGCGTAAAAATGCAAGAAAGGTGCTGAGAGAGATTAAGGCCGACTTCTTCAAACCCGATGTGGATGAAGAAGAACGCGATCGTGCATGGAAGTACATCGAAGCCGGCATTCTGAATATCATGGAAAATTCTGGCGGACGTAATCGATATTGCCGTGAACATAGCAAAGCTTATATTCAGACAGTTCTGAGCGGTAAGGCCGCACCTTTCAAGCGGACTATCAGCGATAGCGACATTATTAGCCGTAGAGAACAGATCGTCTATCAGAAGATTTGGAGATACCTGGAAGCACGTGTTTTTCCGCTGGCCCCGGAAGGGATTGACAGAATTGTAGTGGAACGTACCGCTTTCGACCTGCTGGCTGGTAGCAGAAAGAATATCCAGAATGCCAGCGATCAGTTTGTTGAAGAGATGTACCAACACGGTCCGATGCATGGTTTTTCAAGCACCGCCGAAATGCTGAAAGAAGAGTTCGCAGGTATGTGCGCATATTGTGGCTGTGAAAGCGCCGCCCTGATCGACAGAGATCATATCCTGCCAAGGGCAGATTTCTTCTTCGATTCCTACCTGAATATTGTGCCGGCCTGCCCCAAGTGCAACAGCGACCTGAAGGGTAAGCGGCCCATCTCCACCAGCTTGCTGCGTATCGACGATAAGGCATACGAAGCTTACAGCCATTATCTGCAGAAGATTAGCAAATCTCGGCCCATGCACCTGTTTCATACCATTAAAAAGGGCGTGCTGAATCTGATGAGAGATCCTGAGCGGAGCTGGGAAGTGGAGCGGTACCTGAGTCTGATCGCCAAGCAGTTCAGCGAAATCGTTCAGACACAGAGAAGCCCGCGTCCTTTTGCACGCTACCTGACCAGCAAGCTGATCAAACGTCAGGAGAAGATGCCTGAGATCCTGTTTCGTTCTGGACGTCATATTAATCTGTATCGCACCATCAGCTATCCAGCGTTCAGCAAAGTCGAGGAAAAAGAAGAGGGCAATACCGCCAATCATGCAATTGATGCCATGCTGCTAGCAAGCGATCTGCCGAGTCCGAGCCCGCTGGAAGCACTGAGCATTCCTGTGTCCCTGGTGAAACGCTGGAGCCTCTCCGTTCAAGAAAGCGCCCCTAAGACAGGCCGGAATGGTATTCCGGAGATCCCGTGTCATGGCAGCTACGTGGATGGCTTTGAAAAAGTGGACGGAAATGGTTATGTTGAAATTGAACTGGACAAAATGAGCTGGAACCAGAAGGATCGTATGACCCACAAACAGGACCCTTATGGTTGGAGCCAGAAGGCTCAGATGCCGACCAAACGTACCGCAGCCGTTGATCTGTACAACAGCCTGAAAAAAGAGAGCAACGTGCAGAAAGTGAAGAACATCATCGAACGTATACACCACCCCGCCCTGAAGCGTGTTATGGCCGCTAGCGCGAATAGCGAAAATCCAGGCGGAAGCGTTGCAGAAACCATGAAGAAGTGGCTGAGACAGTCTGTGCAGAACTCTATCAACAACTCCAGCTTTAGCAATCACCCGGGTGATCAGGCACGTAAAGGCGATCTGGAAAAGTTCACCTCGGAGGAGACAGCCATCCCTGCCGTCATCGGCATCAAAATGTTCGACACCGGCGTTAGAGGCAAAATAGATCTGTCCAGAATCGACAAGCAGACCGGCAAAGTTGGCCACAGATATATGACCCAGCCTGCAAACCGGGGTGTGATTCTGGCCTATCCCAAAAAACCTAGCGGTGAACCTGATACCGATAGACCTTATCTGGCCTTCATCAAACAGGATAGCAGCCTGAAGCCGGAGGGTGCAATGTTTAAACCTCTGCCGGAAGGTATTCTGAACGGCAAAATTCTGGGCACCGGCTACTCTCCTGGGGAATGGATGGGCCAGGTTGAAAACTACCTGTCCGAATGCGGTTTTCACAGCTATGTGAGCCTGACCCCTGGTTGTGTTGTTTGTTATAAAAACGGTAAAAAATGGTTTGTGCGCAACTTTGATCAGAGCGCCGATTTCAAAAAAGCAAGACTGAAAGACGTGGCCGGAATCCAACGTACTCCGTTTATCACCCGTATTTCGCCGCTGAAGATGCTGACATAATTAATTAAC**ACCGGT**ATC |
| Cas9d-3 | AGT**GGATCC**ATGGTTAATACACTGGCCATCGACTTCGGCAGCAAATATATTGGTGTTGCTCTGGTCCAGCACAGTGAAACCACCCCTAACCGGGTGCTGTATGCCGCCACCCTGATCGTTGCCGCACGGCCTCTGAAAGCAGCCGTGGAAGATAGAGCTAAAACCCGGAGAATGAGAAGAACCCGTAAGACACATGCTAGAAGACTGCGCCGGCTGGCTCAGGCCCTTGCTGATATTCCGAATGCTGCCCACATTGTGCGTTTTTGCAGACGGCGCGGTTATAGCCATGATCCGGATGATACCGACGAAGGCACAAGAAGCCTGCATTTTCCGCGTGCAGAATTTTTTGAATGTCTGAGACAAGAGGTTGCCCGGATTATCGCACCCGAGGATCAGGAAAGAATGCTGAGAGCATGTGCAAAACATCTTAATGCTGCCAAACGTCGAGAAGCAGAATTGCGGCCGGCCAGGTTCGACAACCGTGGTCCCAGCAAATGTCAGTGGGAAGGATGTCGGCAGAACGTGCCACGTGCAGCCAATGCATTTCGTGAACGTCTGCAACAGGCGCTGTGCGTGTGGATGAAGCCTGTGTTCGATGAGAGCCAGGCTCCTGATCGTTTGCGGCGGAGCGTTGATCATTGGGTTCGGGAACTGGAAGGCCTGGCTCGTTGGTATGCCAAGACCAGAGACTTGGAAGCAGACGCCCAGCAGAGCGAGCGTAAAGCAATTGATAAACGTAGAAAGCTGGTTTATAAAGCCATCAGAGACCGGGTTGCACAGGAGGCCAGCCAGAGAGTTGCAGAAGAGTTCAACGCAAATTGGAGTGAAGGTCGTGGATATCGAGCTAATCTTACGGATATCGTGTGCGGCAAGCAGGGAGGAAGACTGCGTTATTGTCGTAAACATAGCGATTGTTTTGTGGATTTTTTCCTGGCAGGTAAACAGATTCCCAATCGTACAGACATCGCGGAAAGCGATCTATTTGGTAGACGTCAGCAGATTGTCTTTGGCCGTATCTGGCGACTGGTTGAAGATCGTTTACTGCCGCTGGCCGGTGGCGGTATCGATAGAGTGGTGGTTGAACGTGTCGCGTTTGATGTTCTGGCAGGGAAGGCGAAAGACCGGAGAGATCTGAGAGAAGGTGAAGCCGCCGAGATGTACTGGTACGGCCCTCAGCACGGCTTCCAGAGCAGAAAAGAAATGCTGAAAGCAGAATTCGACGGGAGATGCGCCTATTGTGGTCGTCAGGGCGAGTTTCAAGAAGTGGAACATGTTCTGCCTAGAAGCCAGTTTCCGTTTGACAGCTACCTGAATGTTCTGCCGGCATGTAGCGCATGTAATCACCAGAAAGGCAGCCGTACCCCGCTGGCCGCTGGTATGACCATTCATCGTGAGGCCTACAGCGCATATGCAGACTACGTGGGCCGTCTGAAGCCTCCTCATCTGTTTCATACCATCAAGAAAGGTATGCTGAACCTGCTGGCACGAAATGGCAATTGTCACGAGGTGGAAAAGCAACTGGGTATGCTGGCAGATAATCTGGTGACCATTACAACCACCCAGAAAAGCCCGAGACCTCTGGCACGTTATCTGGCCAGTAAACTGGCCAAGACAACCGGCCAACCGTGTGAACCGGGTTGGACCAGCGGCCGCCATACAGCACTGTACCGTGAAATCGTTCTGCCGGATTATAGCAAGCCTGCCGAAAAAGAGCGGGGTGATGTTATTAACCACGCCGTTGATGCAGTGATCCTGGGCTGCAAGTTCCCGTCCGCCGCCGCACTGGAGAACCGTAGATGGTACTTCCGGGCAAGCGATATTAATACCTGGCGAGAGCAGGTGCTGGCAGCCGCGCCTGCACTGGGTAACGGCCTGCCGCGTATTGAACCCGTTGTGTTCTTGCCTTTCTTTGAAGAAGATCTGGGTGGTGGCTACTGCGCCATCGACCTGACCGCCTTCAGCTGGAATCGTGGTCGCCAGAGCGGTCATCTGCTGGACCCTGTGGGAGTTACAGCCCAGGGCCAACGGGTGAAAAGGGACCCAGCCGCCGAAGTGCTGGCCAATCTGAAAAAGGGTTCTGAAAGAGATGAACAGATCCGGAGAATTGCCCATCCGGCCCTGCGCAAGAGACTGCAGGCCGACCCTAACAGAGCCGCTGAGAATCTGGTGCTGTGGCTGCAGGCCAGCGTGCAGGCAGGTCTGGTGAACGGTGGCATGGGCAACCACCCTGGCGACCAGGCAGGTAAACGCCTGCTGGAAGAATTCGTTACCCAGCCGGTGGAGTCTTTTCTGACCGAGGATAAAGATCTGCAGAGGGGCATCCCTCGAGGCATCGGCGTACGTTGCCTGATCGCAGGAGCAGCAGGTAGCTTTGATGTGGCGCGTTGTGATGAAGAGGGTCGCGTTTTTCAGCACTACAAGGCCGACCCTCACTGGCGGTGCCTGTGTGTGGGTTATCGTGAACGAAACGGTGCACTGGATCGTCGTAGGCCCGAGCTGCTGTGGGTGAACCAGGTGCGGGCAGTGTCTCGTGGAGCATCCGGTTCTCAGGTTCCGCTGGATGTCCCCGACGATAGCCCCCTGCGCGGCCGGCCCTTAGGTGGCCGGGGTTCTCTGAAGGCGTTTCTGACCGAATGGCAGCAAGCATTTGATGCCCTGTGCCACGCCGACGGCATTGTTAAAGTTTTTAGAATCACCCAAGGCTGCGTGATCGAAAAAACCGATGGCACCTGTTTTCAGCTGAGAAACTTCATGCGTGAACAGCCGTGGATGAAACGTTCTCCCTTCCGGGACATTCGTCGTGTCTACAGGAGCCCGGTTCGTTTCTGGAGAGCCTGTGCCAACAGCGCCGCTCCGCCAGAGTAATTAATTAAC**ACCGGT**ATC |
| Cas9d-4 | AGT**GGATCC**ATGATTACACTGGGTATTGATTATGGTGCTTCAAATGTGGGTATCGCATTAGTCCGTAATACCGATGAAGGCAACGAGCCACTGTTTGCAGGCACCATGATCCTGGACGCAAGGCTGCTGAAAGAGAAGAGCGAAACCAGAGCAGCGTGGCGTCGTCTGCGCCGTACCCGGAAGACCAAACATCGTAGACTGAGCGATCTGAAACAAAGGCTGCTGAGCCTGGGTTGCGATAGCGATCTGGCAGCAAAAGTTATTAAATTTTGCGAACGTCGTGGCTTTAAATATCGTGATATTGAGAAATTAACCGACGAGGATCTGGCATATGGTATTCCGCGTGAAGAATTTTTTGGCAGCCTTGAAGCCGAACTGGACAGACTGGTGAGCAGCACCCAGCTGCGGGAGAAAGTGCTGAAGGCCTGTGAAGAGGTTCTGAACAGGAAGGGTGATCCCCGCCTGGAAGTTCGTCCTATTCGTATTGATAATAGGGGTGTTAATCGTTGTAGCTGGGAAGGCTGTCCGAATGTTACCCCCAGACGTGAGAACGCAACCGTGGATGCCCTGATGCAGGCACTAGTGAACTACTTCCAAGGTGCGCTGAAAGAAGATCCTGGACTGATCGACCATGTTAAGCAGGCAGTTGATCATCTGAACCGCATTGCTAAGCCGATGCGTGTGGCCGATGATAAGGATGCCAAAACCGAAGTGAATAGCCTGAGACGTGAGGCGAGAAAAATTCTGCGTTCTCTGCGGAGTGAGCTGACCGAGGTGGACCCCACAGATGAGCGGGCAGCGAAAAAATGGAAAGGTGTGGAACAGAACATCGTTAATATGGTGAAATCCGGTGAAGGTCGAAACACCTACTGCAGAGAACATAGCAGCGAATATGTGAGCAGAATCCTGGCAGGTAAGCCGATTGATTTTAAGAAAAGCATCAGCGAGGCTGATATCATCTCACGCCGCGAGCAGATCGCCTTTAGCAAGATCTGGAGATATATTGAAGCCCGACTGCTGCCTCTGGCCCCTGAAGGCATCGATAGAATTGTAGTGGAGCGTACCGCATTCGATCTGATTAACAAGAAAAAATCGCGTCGTGGCGGAAAGAACAAAAAGCCCACCCCTGATCAGGACGGCATGATAGAGGAAATCTACCAGATGGGCCCTATGTATGGTTTCGAGAGCCGCCGGGAAATGCTGACCACCGAATTCGGCGGTCTGTGCGCCTATTGTGGTAAAGCCTCCGGCGAACTGGTTGAATGCGACCATATCCAGCCTCGGAGAGATTTTTTTTTCGACAGCTATCTGAATATCGTGCCGGCATGTCCTCAGTGCAACGCCGAAAAAAGCAAAAGGAGACTGAAGCACACCAGCCTGCACATCAATGCAGAAGCCTATGGTAAGTATGATCTGTACTTGAAGAGCCTGCGTACGAAAGGCCGGCCGCTGCATTTTCTGCATTATGAAAAAAAAGGCATCCTGAATCTCATGAGAAATCCGGAAAGAAACTGGGAAACTGATCGTTATCTGGGCCTGGTGGCCAACAACTTTGCCGCCATCGTGCAGAGCCAGCGTGCACCCCGTCCGTTTGCACGGTTCCTGTATAGCAAAATTAGCGCGCGTCAGGACAAAGATCCGAAGATTGATTTTCGTAGCGGTAGACACACCGCACTGTATCGGACAGTTGCATATCCAGATTTTGACAAGGCCCGAGAAAAATCTATTGGCGGGGCAGTTAACCATGCCCTGGATGCCATGCTGCTGGCATGTCTGCTGCCTGACCTGCACAAGCTGGAAGCTCGAGGCATCAACGTGCACAACCTGGGGACCTGGCGGAGAAGCGTGCTGAGCAAGGCCCCGAAAGCCGCCAAGGATGGTGTGCCTGTGGTTCCTGCACATGGCTGGTGTGTTCCTGGATTCGAGATCCAGGACGAAAACGGTTACGTGAGCCTTGAGATGGCATGTATGAATTGGAACCAGAAAGACAGCGCCACCCACAAGCAGGACCCCTACGGCTGGTCTGAACGCGCGAGGAAACCGACAAAACGTGCAAGCGCAATTGAACTGTACGAGAAATTCGTGAAAGAGAAGAATGAAGGCAAGGTTCGTAACCTGGTGGAAGACATTCATCATCCGGCCCTACGTATGGCCATGACCAAGGCCTTCGAGGCATCCAAAAGTGGTCCGGCCGTTGCCGAGGCCATGAAACAGTGGCTGCAGGTTCAGGTTAAAAATAGCCTGAGTTCCTCTAGCTTTACCAGGCACCCGGCTGACCAGCGTCGTAAACAGGCCCTGGAAAATTTCGCCCAGGGCAGTGATACCAAGATCCCGCAGGTTATTGGTATAAAGAAGTTTGATCGTGGAGTGGTTGGTAAAATCGACCTGGTCCGGGAAGATCCGGAAACCGGCAAACCGGGCCACAGGTATATGGCCCATCCGGCCACCAAAGAAGTGATCCTCGCCTACCCTGCCGCCGCAAATGGTAAGGCAGATCTGCTGAAGCCTTACACCGTTGCAGTGAAGCAGAACTTCAGCCTGCGTACGCAGGGCGAAAGAATTTTCAAACCAAAACCTCAAGAGCTGGAAGAAGGTATAGTGTGGGGCAGAGATGCAAGCCCTACACACGGTTGGGAGGACAAATTTGAAAGATGGCTGCTGGAATGTGGCTTCCACAGCTACACACTCCTGACAAGAGGATGCGTGGTGTGCTACGAAGATGGCACTCAGTGGTTCATCAGAAACTTTGATGATAATAAGGATTTCAAGAAAGCAAAAATCCTGAAAAATGTGGTGGGCGTTAAAAGAACACCATTCGCCAGCAGAGTCGCTAGACTGAAGATCCTGACCAGTGGTAGCCAGAAGCCGTAATTAATTAAC**ACCGGT**ATC |
| Cas9d-5 | AGT**GGATCC**ATGCCTAGCACCACCACGCTCGCAGTGGACTACGGCTCCAAAAACATTGGTATTGCACTAGTTCGAAACCAGGATGATCACAACGTACCGCTGTTTGCAGGAACCGCCCTGTATGATCGTTTTACCCTGTCAAAAAAAGCAGAGCCAAGAGTGCAGATGAGAAGAATGCGGCGGACCAAGAAGAGCAAAAAGGCACGTTTACGGTGGCTGGAAGCAGGTCTGCTGACCCTGGGCCTGGACCACGAGACAGTTCGTCATCTGGTGAACTTTTGCCGTCGTAGAGGATATAAGTCCCTGTTCGATGAAGGCAAAGAAGAGACCAGGGGTAAAAAAGATCAGGAAGAGGACGAAGTGTTCAGATTCAGCCGGGAAGAGTTCTTTCTGGCGCTGGAAAAGCTGCTGGAAACCACACTGCCGGAAGAACAGCTGGCCCCTATCCTGACTCTGTGTGAACACGTGCTGAATCGTCATGGTGATAGATACAAAGAAGTCCGCCTGATTCGTATTGATAATCGGGGTGTGTCCCGTTGCGCCTGGAACGGCTGCAACAGCGTGACCCCGAGACGTGACAACGCCCTGCGCGATGCTCTCGCACAGTTCGTGTGTACCGTTTATGCAAGCGACCTGCGTGGTAATCCTGATCTGTATCAGGAAATCCAGGTGATGCTGGACAAGGTTTCTGAACTGGGCAAGCGATTGCGGAATGCAGGCGGCAGTGACCCCGCCAAAGAACGTAAAGTTCTGCGTAAACAGATTGGCCAGAGACTGAAGAAACTGAAGGGACTGAGCGGTCTGGCTGAGGAGTTACCCGACGAGGCCGTTGGTCAAGGTGAAAAATGGGTGAGCATCCGTCGTAATCTGATGGACCTGATTGAGAGGTCTCGTGGAAGAAATAGGTTCTGCAGACAGCATAGCGACCAGTACTGTAGCCACCTGCTGGCAGGTCGTCCGATTCCGTTCAAAAAAAGCCTGACCGAGGCTGACATCACCAGCAGACGGGAGGAAATTCTGTTTCAGAAACTGTGGCGGTATATCGAGGCCAGAGTTCTGCCTCTGGTGCCACAGGGCGTGGAACGCGTGGTGGTTGAGAGAACCGCATTTGATCTGCTAGCAGGTACTCGGAAGCAGCGCCTGGCCATCAGCGATAAAGAACTGGAAGCCATGTATCAGGAAGGTCCTAGACACGGCTTCAGGAATGATCTGGAAATGCTGAAAGCAGAATTCGGTGGACTGTGCGCCTATTGTGGCAAACCGTGTGGCGATCTCCTGGAACGAGAACATATACTGCCCCAGGCCAACTTCTTTTTCGACAGCTACCTGAACAAGGTGCCCGCATGTCCGGATTGTAATCGTATGTGGAAGGGCAAGGCAAGCCCTCTCGCAGCTGGTCTCGTTGTTCATGAAGAAGCCTTTAAAGCATATAGCAAATATCTGGATACCAAATTTAAAGCCAAGAATAAGCCACCGCATGCATTTCATACCATCAAGAAAGGCATCCTGAAGCTGATGACCCAGCGTGATCGTCTGCCAGAGGCCGAAATGCTGCTGGCCATCATCGCCGATAACCTGGGAGCTATTGTGGAAGCCCAACGTGGACCTAGACCTCTTGCACGTTATCTGTGTGAAAAACTGAGACAGAGATACGGTCAAGTGCCTCAGGTTGCCTTTCGTAGCGGCCGTCATACAGCAATCTGGCGTCAGGCTGCTTACCCCGATTTTAACAAAGCCGCAGAAAAAGTGGCCGGCGGCACCCTGAACCACGCCCTGGATGCGCTGATCATGGCCTGCAAGCTGCCTGATATCACCGTTCTGGAGGCAAAAAACCTGGCCAGCTGGACCCTGGCGTCTTGGGCAGAAAAGGTCCAGGCCGCAGCACCGCCTCCGGGCGAAGATGGTGTTCCGGTTCTGCCTGTGCCCAGCTTTGCCGTGCCAGGTTTTGAGGAAGTTCTGCCTGGTAATTATGTTCAGACCGATCTGGCCCTGCTGAATTGGAATCGTAAAGATAGCAGAGTCCAACGCCAGGACGCCTACGGCTGGGATGATTGGGAAGATATGCCGGCGAAGCGGGTTAGCGCCGCGGAGCTGGCAGCCCTGCTGCGGGACGCTGACAAGAAACAGACACCGGCCGAACGTCAAGCAGAGGTTCGTAAAGAAGTTGAAGTTATTGTTCATCCTAGACTGCGGCAGTCCCTGGTTAAAGCCAATACCGGCGAAAAACCGGGTGAAGCAACAGCGCAGGCACTGACCACCTGGCTGAGGAAGACCGTGAAGAGAAGCCTGAAAAGCGCAAATTTTTCTAGCCATCCGGCAGACCAGGCTCGTGGCAAGGCCCTTCGTGATTTCGCTCAGGGCGTTACCGACGCACTGCCGGCAGTGATTGGTGTGCGGATTCGGTATCCTTGGCTGAGAGCCAACATGGATCTGAGCCGGGTGGATCGCCATACCGGCGCCCTAGTGCACAGCTACGTGGCTGATCCGGCAAATCAGGGTATCATTGTTGCTTACGCCGAACGTGATGGCAAGATGCACAGAGATAAACCTCTGACACTGGAGCTGAGACAGAGCGGAGCAGTTATCCCTGGCGTGAGAGCTCTGGGCGAAGTGCCGAGCGGCCCTCTGTGCGGTCGTGCGCTCGGCCAACCGAAGATCAACCCCGAAGAATGGTCCAAGGCCTTACGTTCTTACCTGCGGGGCGCCGGAATTAAAGAATATACCATGGTGACCCAGGGTTGTGTGGTCCACTACGAGGACGGTAGCAAAAAGTACATCAGGAATTTTAGCAGTAGCTATGGCTTTAAAAAAAGCCTGCTGAAAGGTATCATCGGTATGCGGCACAGTCCGCTGGCCAGCAAAGTGACACCTAATGCCCGAATCTAATTAATTAAC**ACCGGT**ATC |
| MG102-2 | AGT**GGATCC**ATGGTTACACTGGGTGTGGATTATGGTGCAAGCAGCGTTGGACTCGCGCTGGTGAGAAGCGACGAGAAAGGGCATAACATTCCCCTGTTCGCAGGCACTATCAGACTGGATGCTAGATGGCTGAAGGAGAAAGTTGAGGTGCGTGCAGGAATCCGTCGTCTCAGACGTACCAAGAAGACGAAGCGCCACAGACTACGTCAGCTGGAACAGAGCCTGACCCAGATTGGCCTGGGCGCAGAGCAGATTCAGAGCATTGTGCGGTTTTCCAACCGTCGTGGTTATAAAAGCCTGTTTGATGCCGGTGTGGCCGACGATGATCACGATGAAGCAGAACTAACCTACCGATTCACCCGGGAAGAGTTCTTCAAAGCCCTCGGCACAGAACTGCAGTCTATCATCCCGGATACTAGCCAGCGTCAGAAGGCACTGAGCACCTGCGAAACAATCCTGAACAGACAGGGCGACCGCTCTCTGGAAATCAGACAGATCAAAATTGACAATCGGGGTGCAAGTCGGTGCGCATGGGAGGGTTGTAATCGTGTGACCCCTCGTTCCGACAACGCACTAGGCGAAGTGCTGAGCCAGCAGATCTACACAGTTTTTCAGAGCGCCCTGAAGGTTAATCCTGTGTTGTGCCAGAAAGTGGAGACAGCCACCCATGAACTGCATGAACTGGCCAAACGGCTGCGTAATGCATCAGGCGAGGCCGCAAGCAACGAAAAGAAAATTCTGAGAAAACGTGCCAGAACCATTCTCCGAGCTCTGAAAGAACTGCTGTATACCCCCGCCGAAGGCGTTGGCGATGCTGAAAAAGCCTGGAAATACATCGAAACCGGCATCATGAACATCATGGAAAGTCGCCTTGGAAGAAACCGTTATTGCAGAGAACATAGCCGTGAGTACGTGCAGACCATTTGCAGCGGCAAACTGGTGCCTTTTAAGCAGACCATTAGCGAAAGCGATATTATTAGCCGTCGTGAACAGATCGCCTACGCCAAGCTGTGGCGTTATATTGAAGCACGCATCCTGCCTCTGACCCCGGGTGGTATCGACCGTATCGTTGTCGAACGTACCGCCTTTGACCTGCTGGCCGGTAGCCGAAAAACCATCCAAAAGGCAACCGATCAGTTCAAGGAAGAAATGTACCAGCACGGTCCGATGTACGGCTTTGAAAGCGTTCCGGACATGCTGAAAGAGGAATTTGGTGGCCTGTGTGCCTATTGTGGCCAATCCTCTAGCACCCTGATAGACGTGGACCATATCTTACCGCATGCCGATTTTCTGTTTGATAGCTACCTGAATATTCTGCCCGCCTGTCCGAAGTGCAACAGCGACCTGAAGGGTGATCGTAGCGTGAGCGATGCCAGCCTGACCATCCATCCCGATGCGTATCGTGCCTACAGCGACTACCTGAAAAAGAAATTTAGCACCAGACCGATGCACTACTTCCATTCGATCAAGAAGGGAGTGCTGAATCTGATGCAGGACGCCTCAAGACTGTGGGAGGCTGAACGTTACCTAAGCCTGATTGCTCGTCAGTTCGGCCAGATTGTGCAGAGCCAGAGAGGACCTAGGCCGTTTGCACGGTATCTGAGCACCAAACTGAGTCGTCGTCAGGGTCAGGCACCGGCCATTCGCTTCAGAAACGGCAGACACACCGCACTGTACAGACGGGTTGCATATCCAGATTTCCAGAAACAGGCAGAAAAAGCCGAAGGCAACGTGCTGAACCACGCCCTGGATGCAATTCTGCTGGCCAGCGAACTGCCCGATCTGTACCCGGTGGAAGCCCTGAATCTGCCGCTGTGGCAGCTGAAAAATTGGGCAGATACCGTTCGGGCCCGAGCTCCTAAGAGCGGTGATGAAGGTGTCCCCATTTGTCCGGATGGCCCTCAGTACGTTGATGGCTTCGAGACCGTGCATCCCGGTGGTTATGTTGAAGTTGATCTGAGAAGCATGCGTTGGAATCAGAAGGATTCCATGACCCACAAGCAGGACCCTTATGGCTTCAGCGAAAAAACCGGCATGCCTACCAAACGGGGTAGCGCATTTGACCTGTATACCAAGCTGAAGAAAGAAAAAAATCCGAGCAAAGTGAAAAGCCGTATTGCCCTGATCCACCACCCTGCACTGCGGAAAGCTCTGAGCGAGTCTCTGCAATCGGATACTACAGGCAGCTCTGCCGCCGAAAACCTGAAGACCTGGCTGAGAATCAGCGTTAAAAATAGCCTAGCTCACAGCAGGTTTAGCAATCACCCTGGCGACCAGGCACGTCGGAGCGAGCTGGAAAAGTTCATCACCCAAGAGGACTGCCCTATCCCTAGCGTGATCGGCGTGAAGATGTTTGATATGGGCGTGAGGGGTAAAATCGACATGAAGCGGCTGGACCGGCAGACAGGTGGCATCGGACATCGTTACATGACACAGCCTCCGAATAAAGGCGTGATTGTTGCATATCCGAAAAGAGAAGATGGGAAACCTGACCTGACCAAACCGTGTTGTGTTTATCACAAGCAGGACCTGAGCATGACCCCTGAGAACATCAGCATCTTTAAACCGCCCCCACCGATCCTATCTGATGGTGTAATCCTGGGCAAGAAACCGTATGATAAAGGTGATCGTAGAAAGGCTCTGGAAAAATACCTGAGCGATTGTGGTTTCCATAGTTATGTTTATCTGACGCCTGGTTGTACCATTCGGTACAAAGACGGTAACGAGTGGTTCGTGCGTAATTTCGACAGCAGCGAAGATTTTAAAAAAGGACGTCTGCGCGAGATCATCGGCACACGGAGAACCCCTTTTGTTGACTCTCTGATCCCACTGAAGGTTCTGTCCCAGTAATTAATTAAC**ACCGGT**ATC |

Yellow highlighted: BamHI restriction site

Pink highlighted: AgeI restriction site

**Table S2.** **sgRNA expression templates designed for cloning into a T7 promoter–driven vector for *in vitro* transcription**

| **Name** | **Sequence (5’---3’)** |
| --- | --- |
| Cas9d-1 sgRNA scaffold | CACCGCTAGCTAATACGACTCACTATAGGGTCTTCGAGAAGACCTGTTTCAATTAAACTGATTGAAAAATCAGTTCCGGTTGAAAAGAGCATCCGCCGGACAGCGCACTCCGGGATCGGGCAGTCCCGGCACTTGCAGTTTCCCCGGTTAGTCCCTCGGAAACAAGCCGTCCGGCATGTCGAAAGACAGGATGTGAGCCCATTAAAGCTTGGAT |
| Cas9d-2 sgRNA scaffold | CACCGCTAGCTAATACGACTCACTATAGGGTCTTCGAGAAGACCTGTTTCAATCAAACTGATTGAAAAATCAGTTCCGGTTGAAAAGAGCATCCGTCTGAAGGGCACTCCGGGATAGGGCAGTCCCGGCTCTTGCTGTTTCCCTGGCACATCACTGTGCTCCGGAAATGAACCCTCGGACAAGTCGAAAGACAGGATGTGAGCCTAATTTAAAGCTTGGAT |
| Cas9d-3 sgRNA scaffold | CACCGCTAGCTAATACGACTCACTATAGGGTCTTCGAGAAGACCTGTTTCAGCCAACCCAGAAATGGGTGGTGACTGAAAAGAGCCACGCGGCCGGAAGCCTGCACCCTGGAATTGGACAGTTCCAGGCTCTGCGCGACGGGTACAGAGAAGCCCGTCGACTGGCACCGGCCAATGTGGTGAGTCCATGTTTTTTAAAGCTTGGAT |
| Cas9d-4 sgRNA scaffold | CACCGCTAGCTAATACGACTCACTATAGGGTCTTCGAGAAGACCTGTTTCAATCAAACTGATTGAAAAATCAGTTCAGGTTGAAAAGAGCATCTGTCTGGCGAGCACTCCGGGATGGGGCAGTCCCGGCTCTTGCGGTTCACGACTCAGCTTTGGGTCGTTGGAACACATCGTCAGACACGTCGTAAGACAAGATGTGAGCCCGTATTTAAAGCTTGGAT |
| Cas9d-5 sgRNA scaffold | CACCGCTAGCTAATACGACTCACTATAGGGTCTTCGAGAAGACCTGTTTCAGTCACCCTGAACGAGAAATCGTTCAGGTTCAGGCTGGAAAGAGCATCTGTCCGGGAGGCCACTCCGGGTCAGGGCAGACCCGGCTCTTGGTGCCTCTTTGGCTCCGGCTCTGGAGGAAACCTTCCGGCATGTCCACTGGACCAGATGTGAGCCCGATTTAAAGCTTGGAT |
| MG102-2 sgRNA scaffold | CACCGCTAGCTAATACGACTCACTATAGGGCACGGGCAGCTTGCCGGGTTTCAATCAAACTGAAAAGTTCCGGTTGAAAAGAGCATCCGTCCGGAGGGTGCACTCCGGGATGGGGCAGTCCCGGCACTTGCGTTTTCCCCGGCTTACGCTTCGGAAAAAGGCCCTTCGGCACGTCGAAAGACAGGATGTGAGCCCAATTTTAAAAAGCTTGGAT |

Yellow highlighted: T7 promoter

Pink highlighted: NheI restriction sites

Red hilighted: BbsI restriction sites

Green highlighted: HindIII restriction sites

Underline: Spacers

**Table S3. Oligonucleotides and primers used for construction of *in vitro* sgRNA templates**

| **Primer Name** | **Sequence (5’---3’)** | **Used for** |
| --- | --- | --- |
| T1_20nt_vitro_TOP | TATAGGGCACGGGCAGCTTGCCGG | Top oligonucleotide used for cloning 20-nt spacer into BbsI-digested sgRNA templates |
| GTTT_T1_20nt_vitro_BOTTOM | AAACCCGGCAAGCTGCCCGTGCCC | Bottom oligonucleotide used for cloning 20-nt spacer into BbsI-digested sgRNA templates |
| RTW443_T7_F | GCTAGCTAATACGACTCACTATAG | PCR amplification of T7-driven sgRNA expression templates, forward primer |
| sgRNA_Cas9d-1_R | TTAATGGGCTCACATCCTGTC | PCR amplification of Cas9d-1 sgRNA expression templates, reverse primer |
| sgRNA_Cas9d-2_R | ATTAGGCTCACATCCTGTCTTTC | PCR amplification of Cas9d-2 sgRNA expression templates, reverse primer |
| sgRNA_Cas9d-3_R | AACATGGACTCACCACATTGG | PCR amplification of Cas9d-3 sgRNA expression templates, reverse primer |
| sgRNA_Cas9d-4_R | AATACGGGCTCACATCTTGTC | PCR amplification of Cas9d-4 sgRNA expression templates, reverse primer |
| sgRNA_Cas9d-5_R | AATCGGGCTCACATCTGGTC | PCR amplification of Cas9d-5 sgRNA expression templates, reverse primer |
| MG102-2_Cas9d-5_R | ACTGATAATTGGGCTCACATCCTG | PCR amplification of MG102-2 sgRNA expression templates, reverse primer |

**Table S4.** **Complementary oligonucleotide pairs used for cloning target sequences with variable PAMs into the pUC19 backbone**

| **Oligonucleotide Name** | **Sequence (5’---3’)** | **PAM (5’---3’)** |
| --- | --- | --- |
| PAM30_Top | AATTCGGGAGGGGCACGGGCAGCTTGCCGGCACGAAAAATGCGG | CAC |
| PAM30_Bottom | GATCCCGCATTTTTCGTGCCGGCAAGCTGCCCGTGCCCCTCCCG |  |
| PAM31_Top | AATTCGGGAGGGGCACGGGCAGCTTGCCGGTACCTTAAATGCGG | TAC |
| PAM31_Bottom | GATCCCGCATTTAAGGTACCGGCAAGCTGCCCGTGCCCCTCCCG |  |
| PAM32_Top | AATTCGGGAGGGGCACGGGCAGCTTGCCGGGGCACTAAATGCGG | GGC |
| PAM32_Bottom | GATCCCGCATTTAGTGCCCCGGCAAGCTGCCCGTGCCCCTCCCG |  |
| PAM33_Top | AATTCGGGAGGGGCACGGGCAGCTTGCCGGAGCGTAAAATGCGG | AGC |
| PAM33_Bottom | GATCCCGCATTTTACGCTCCGGCAAGCTGCCCGTGCCCCTCCCG |  |

**Table S5. Adapters and PCR primers used to enrich cleaved and uncleaved dsDNA fragments in *in vitro* PAM characterization for deep sequencing**

| **Primer Name** | **Sequence (5’---3’)** | **Used for** |
| --- | --- | --- |
| TK-117 | CGGCATTCCTGCTGAACCGCTCTTCCGATCT | Generation of an adapter with a 3’ dT overhang for capturing cleaved dsDNA fragments, adapted from Karvelis et al. ^14^ |
| TK-111 | GATCGGAAGAGCGGTTCAGCAGGAATGCCG | Generation of an adapter with a 3’ dT overhang for capturing cleaved dsDNA fragments, adapted from Karvelis et al. ^14^ |
| Library_RTW554_Cut_F1 | GGCGTTTCACTTCTGAGTTCGGC | PCR amplification of cleaved dsDNA fragments |
| Library_RTW554_Cut_F2 | CAGACCGCTTCTGCGTTCTG | PCR amplification of cleaved dsDNA fragments |
| Library_RTW554_Cut_F3 | TCGCAACGTTCAAATCCGCTCC | PCR amplification of cleaved dsDNA fragments |
| Library_RTW554_UC_R1 | GCCTGACTCACTATAGGGAGACCG | PCR amplification of uncleaved dsDNA fragments |
| Library_RTW554_UC_R2 | CGGTATTTCACACCGCATACGTACG | PCR amplification of uncleaved dsDNA fragments |
| F1a_library | TCGTCGGCAGCGTCAGATGTGTATAAGAGACAGAAAACACACCGCATACGTACGATTTA | Addition of Illumina adapters to uncleaved samples, adapted from Pedrazzoli et al. ^20^ |
| F4a_invitro | TCGTCGGCAGCGTCAGATGTGTATAAGAGACAGCTGCTGAACCGCTCTTCCGATC | Addition of Illumina adapters to cleaved samples, adapted from Pedrazzoli et al. ^20^ |
| F5a_library | TCGTCGGCAGCGTCAGATGTGTATAAGAGACAGCGTACGATTTAAATAGGCCTGACT | Addition of Illumina adapters to uncleaved samples |
| R1_library | GTCTCGTGGGCTCGGAGATGTGTATAAGAGACAGCGTTCTGATTTAATCTGTATCAGGC | Addition of Illumina adapters to cleaved samples, adapted from Pedrazzoli et al. ^20^ |
| R2_library_Cut | GTCTCGTGGGCTCGGAGATGTGTATAAGAGACAGTCCTACTCAGGAGAGCGTTCAC | Addition of Illumina adapters to cleaved samples |
| R3_library_UC | GTCTCGTGGGCTCGGAGATGTGTATAAGAGACAGCAGTCTTTCGACTGAGCCTTTCG | Addition of Illumina adapters to uncleaved samples |

**Table S6.** **Complementary oligonucleotide pairs used for guide RNA cloning into BbsI-digested sgRNA backbone vectors**

| **Oligonucleotide Name** | **Sequence (5’---3’)** |
| --- | --- |
| *CLTA* target-1 NAC_Top | CACCGTCTACAAACAACCCTTCGCT |
| *CLTA* target-1 NAC_Bottom | AAACAGCGAAGGGTTGTTTGTAGAC |
| *CLTA* target-2 NAC_Top | CACCGGAAGTATGAGCAAAGGTAAA |
| *CLTA* target-2 NAC_Bottom | AAACTTTACCTTTGCTCATACTTCC |
| *HBB* target-3_NAC_Top | CACCGTTCCCTAAGTCCAACTACTA |
| *HBB* target-3_NAC_Bottom | AAACTAGTAGTTGGACTTAGGGAAC |
| *HBB* target-1_NAC_Top | CACCGACTTTCTTGCCATGAGCCTT |
| *HBB* target-1_NAC_Bottom | AAACAAGGCTCATGGCAAGAAAGTC |
| *AIFM* target_NAC_Top | CACCGTGTGTGACTGTACCAGGTAG |
| *AIFM* target_NAC_Bottom | AAACCTACCTGGTACAGTCACACAC |
| *AIFM* target-2_NAC_Top | CACCGCAACTTAGATAACACCTAGT |
| *AIFM* target-2_NAC_Bottom | AAACACTAGGTGTTATCTAAGTTGC |
| *Casp3* target_NAC_Top | CACCGAGAAACTCCTCCCAAACCTA |
| *Casp3* target_NAC_Bottom | AAACTAGGTTTGGGAGGAGTTTCTC |
| *Casp3* target-2_NAC_Top | CACCGACTGTACATAACCCTAATCT |
| *Casp3* target-2_NAC_Bottom | AAACAGATTAGGGTTATGTACAGTC |
| *EMX* target_NGC_Top | CACCGAACTCGTAGAGTCCCATGTC |
| *EMX* target_NGC_Bottom | AAACGACATGGGACTCTACGAGTTC |
| *EMX*_target-2_NGC_Top | CACCGAGTCCGAGCAGAAGAAGAAG |
| *EMX*_target-2_NGC_Bottom | AAACCTTCTTCTTCTGCTCGGACTC |
| *CLTA*_T1_NAC_24nt_Top | CACCGGCTTTCTACAAACAACCCTTCGCT |
| *CLTA*_T1_NAC_24nt_Bottom | AAACAGCGAAGGGTTGTTTGTAGAAAGCC |
| *CLTA*_T2_NAC_24nt_Top | CACCGAGTTGAAGTATGAGCAAAGGTAAA |
| *CLTA*_T2_NAC_24nt_Bottom | AAACTTTACCTTTGCTCATACTTCAACTC |
| *HBB*_T3_NAC_24nt_Top | CACCGTTTGTTCCCTAAGTCCAACTACTA |
| *HBB*_T3_NAC_24nt_Bottom | AAACTAGTAGTTGGACTTAGGGAACAAAC |
| *HBB*_T1_NAC_24nt_Top | CACCGGAGCACTTTCTTGCCATGAGCCTT |
| *HBB*_T1_NAC_24nt_Bottom | AAACAAGGCTCATGGCAAGAAAGTGCTCC |
| *AIFM*_T1_NAC_24nt_Top | CACCGTATGTGTGTGACTGTACCAGGTAG |
| *AIFM*_T1_NAC_24nt_Bottom | AAACCTACCTGGTACAGTCACACACATAC |
| *AIFM*_T2_NAC_24nt_Top | CACCGGAACCAACTTAGATAACACCTAGT |
| *AIFM*_T2_NAC_24nt_Bottom | AAACACTAGGTGTTATCTAAGTTGGTTCC |
| *Casp3*_T1_NAC_24nt_Top | CACCGGTCAAGAAACTCCTCCCAAACCTA |
| *Casp3*_T1_NAC_24nt_Bottom | AAACTAGGTTTGGGAGGAGTTTCTTGACC |
| *Casp3*_T2_NAC_24nt_Top | CACCGTTATACTGTACATAACCCTAATCT |
| *Casp3*_T2_NAC_24nt_Bottom | AAACAGATTAGGGTTATGTACAGTATAAC |
| *EMX*_T1_NGC_24nt_Top | CACCGTAGAAACTCGTAGAGTCCCATGTC |
| *EMX*_T1_NGC_24nt_Bottom | AAACGACATGGGACTCTACGAGTTTCTAC |
| *EMX*_T2_NGC_24nt_Top | CACCGCCTGAGTCCGAGCAGAAGAAGAAG |
| *EMX*_T2_NGC_24nt_Bottom | AAACCTTCTTCTTCTGCTCGGACTCAGGC |

**Table S7. Backbone plasmid sequences for cloning guides**

| **Name** | **Sequence (5’---3’)** |
| --- | --- |
| Cas9d-1 sgRNA backbone | CTGGATCCGGTACCAAGGTCGGGCAGGAAGAGGGCCTATTTCCCATGATTCCTTCATATTTGCATATACGATACAAGGCTGTTAGAGAGATAATTGGAATTAATTTGACTGTAAACACAAAGATATTAGTACAAAATACGTGACGTAGAAAGTAATAATTTCTTGGGTAGTTTGCAGTTTTAAAATTATGTTTTAAAATGGACTATCATATGCTTACCGTAACTTGAAAGTATTTCGATTTCTTGGCTTTATATATCTTGTGGAAAGGACGAAACACCGGGTCTTCGAGAAGACCTGTTTCAATTAAACTGATTGAAAAATCAGTTCCGGTTGAAAAGAGCATCCGCCGGACAGCGCACTCCGGGATCGGGCAGTCCCGGCACTTGCAGTTTCCCCGGTTAGTCCCTCGGAAACAAGCCGTCCGGCATGTCGAAAGACAGGATGTGAGCCCATTAATTTTTGGCCGGCATGGTCCCAGCCTCCTCGCTGGCGCCGGCTGGGCAACATGCTTCGGCATGGCGAATGGGACTTTTTTTTAAGCTTGGG |
| Cas9d-4 sgRNA backbone | CTGGATCCGGTACCAAGGTCGGGCAGGAAGAGGGCCTATTTCCCATGATTCCTTCATATTTGCATATACGATACAAGGCTGTTAGAGAGATAATTGGAATTAATTTGACTGTAAACACAAAGATATTAGTACAAAATACGTGACGTAGAAAGTAATAATTTCTTGGGTAGTTTGCAGTTTTAAAATTATGTTTTAAAATGGACTATCATATGCTTACCGTAACTTGAAAGTATTTCGATTTCTTGGCTTTATATATCTTGTGGAAAGGACGAAACACCGGGTCTTCGAGAAGACCTGTTTCAATCAAACTGATTGAAAAATCAGTTCAGGTTGAAAAGAGCATCTGTCTGGCGAGCACTCCGGGATGGGGCAGTCCCGGCTCTTGCGGTTCACGACTCAGCTTTGGGTCGTTGGAACACATCGTCAGACACGTCGTAAGACAAGATGTGAGCCCGTATTTTTGGCCGGCATGGTCCCAGCCTCCTCGCTGGCGCCGGCTGGGCAACATGCTTCGGCATGGCGAATGGGACTTTTTTTTAAGCTTGGG |
| MG102-2 sgRNA backbone | CTGGATCCGGTACCAAGGTCGGGCAGGAAGAGGGCCTATTTCCCATGATTCCTTCATATTTGCATATACGATACAAGGCTGTTAGAGAGATAATTGGAATTAATTTGACTGTAAACACAAAGATATTAGTACAAAATACGTGACGTAGAAAGTAATAATTTCTTGGGTAGTTTGCAGTTTTAAAATTATGTTTTAAAATGGACTATCATATGCTTACCGTAACTTGAAAGTATTTCGATTTCTTGGCTTTATATATCTTGTGGAAAGGACGAAACACCGGGTCTTCGAGAAGACCTGTTTCAATCAAACTGAAAAGTTCCGGTTGAAAAGAGCATCCGTCCGGAGGGTGCACTCCGGGATGGGGCAGTCCCGGCACTTGCGTTTTCCCCGGCTTACGCTTCGGAAAAAGGCCCTTCGGCACGTCGAAAGACAGGATGTGAGCCCAATTTTTGGCCGGCATGGTCCCAGCCTCCTCGCTGGCGCCGGCTGGGCAACATGCTTCGGCATGGCGAATGGGACTTTTTTTTAAGCTTGGG |

Underline: Human U6 promoter

Yellow highlighted: gRNA Scaffold

Red highlighted: BbsI restriction sites

**Table S8. Primers used for PCR amplification of U6-sgRNA cassettes from positive clones and for the Sanger sequencing of expression constructs**

| **Primer Name** | **Sequence (5’---3’)** | **Used for** |
| --- | --- | --- |
| U6 gRNA-seq-F | CGGTACCAAGGTCGGGCAGG | PCR amplification of U6-sgRNA cassettes, Sanger sequencing |
| HDV gRNA-seq-R | GAGGTACCTCGAGCGGCCC | PCR amplification of U6-sgRNA cassettes, Sanger sequencing |
| T7-F | TAATACGACTCACTATAGGG | Sanger sequencing |
| CMV-F | CGCAAATGGGCGGTAGGCGTG | Sanger sequencing |
| BGH-rev | TAGAAGGCACAGTCGAGG | Sanger sequencing |

**Table S9.** **Genomic on-target sequences**

| **Name** | **Ampicon sequence (5’---3’)** |
| --- | --- |
| *CLTA* target-1 | GCTCCTCAGAGCACAGTTGTACCTCAATTGTGGATTTTAGATGTTTCTGCTTCTCAATGTTCTCTCTTTTTTCCTGCCTGCTTGCCTGCCTTTTGGACCTCTTGCTGTCTAGGGTGGCAGATGAAGCTTTCTACAAACAACCCTTCGCTGACGTGATTGGTTATGTGTATGTTGCAAAACTAATTCCTTTTTCTACATCTGATTCTTTCTACTTTTGTTTAGAGTTAATGCTCCTTTTATGTCACAAGTTGCTTTGCTTTTTAAATTATTTCAAATTGGCACTTTGGGGGCTGCCTAAGAATTGATAAGCGGGGTATGATCTGTTGATGAATCTTCCAGATTTGTGACTCCCTGCATCC |
| *CLTA* target-2 | TTTGGTAATCACTGTGTGCTATTTATCGACTATAGATTTTAGGCTTCAACTGGTTATTAACCTCATTCACTCACTTGTTCACAAAATATATGCTCAGTACTGATATATGCATACCTTATAACAACTCACGTATGTATAAAGCCTAGTCAGTTGAAGTATGAGCAAAGGTAAATACCTGTTTCTCTTCCAATCTGACGTGTTCCCAGGTTGATCACAGGTGTTCCAAAAGCTCCAACTTCTCGAACCCCACAGCTGCTGTTCCCAATCATACAGCCAGCATGGGCAACCAACTGTATAAACTGGTCAAATGGGACGTGTTTAACTGCACGAAAGTTGGGATGATGCTCAATGCCCTTCTTCCGCATCACTCGAACCAT |
| *HBB* target-1 | GCAATCATTCGTCTGTTTCCCATTCTAAACTGTACCCTGTTACTTCTCCCCTTCCTATGACATGAACTTAACCATAGAAAAGAAGGGGAAAGAAAACATCAAGGGTCCCATAGACTCACCCTGAAGTTCTCAGGATCCACGTGCAGCTTGTCACAGTGCAGCTCACTCAGTGTGGCAAAGGTGCCCTTGAGGTTGTCCAGGTGAGCCAGGCCATCACTAAAGGCACCGAGCACTTTCTTGCCATGAGCCTTCACCTTAGGGTTGCCCATAACAGCATCAGGAGTGGACAGATCCCCAAAGGACTCAAAGAACCTCTGGGTCCAAGGGTAGACCACCAGCAGCCTAAGGGTGGGAAAATAGACCAATAGGCAGAGAGAGTCAGTGCCTA |
| *HBB* target-3 | CTGACCTCCCACATTCCCTTTTTAGTAAAATATTCAGAAATAATTTAAATACATCATTGCAATGAAAATAAATGTTTTTTATTAGGCAGAATCCAGATGCTCAAGGCCCTTCATAATATCCCCCAGTTTAGTAGTTGGACTTAGGGAACAAAGGAACCTTTAATAGAAATTGGACAGCAAGAAAGCGAGCTTAGTGATACTTGTGGGCCAGGGCATTAGCCACACCAGCCACCACTTTCTGATAGGCAGCCTGCACTGGTGGGGTGAATTCTTTGCCAAAGTGATGGGCCAGCACACAGACCAGCACGTTGCCCAGGAGCTGTGGGAGGAAGATAAGAGGTATGAACATGATTAGCAAAAGGGCCTAGCTTGGACTCAGAATAATCCAGCCTTATCCCAACCAT |
| *AIFM* target-1 | GGCTGACAACTTCTTATTGACCTGGAGTTTTGGGGTGGTGATGGAAATGTTCTGAAATCAGATAGTGGTGATGGTGGCACAACCTTGTGAATATACTGAAATCCACAGAATTCCACACTTTAAAATGGTGAATTTGATGTGAATTATATCTCAATAAAACTTTTTCAATGCTAATTCATCTCTACCTCTTTTGTGTATGTGTGTGACTGTACCAGGTAGTACAGCTGGATGTGAGAGACAACATGGTGAAACTTAATGATGGCTCTCAAATAACCTATGAAAAGTGCTTGATTGCAACAGGTGAGCATTTCTGGAGATGGCTTTCTTCCCTGATACAGCTCATTCAAGTTTTAGAGTCCAGCCTTCTCTATGGAAGG |
| *AIFM* target-2 | CGATCAGCTTAGCAGGTCACAGAGGAATTATTTTGTGGGTGTTTTTTTCCTAACTTGGCAGGGTTTACTGTGACTAAGCCGATCAGTCTAAAGTTGGATCCTGTAAGGTTGAGTGAACCAACTTAGATAACACCTAGTAACACCCTTGATTGAAGGAAGCTGTTTGTTTTACTTAAGCCATTACAAGTTTAGGGGTTTTTTTCTACATTGGTAAGACAATGAATTTATCTTTGTTCCTTGGTTGCTAGCTTAAAGAAAGGGACATGAAAATTATTTGAAGCTACCTATATCCTCATCCCCTGATGTGACTCCCAAACGGACA |
| *Casp-3* target-1 | AGGCCTAGTAGGGTGTGTGACCCATAGTTGGAAACCTAGTCCTGGAACCAAAAGGACACCACAGGAGAGGCAGGACTCCCAAGCGCCGGTGCTTTCTACCCTCCCGAATCGCTCTATAGGATTAGAGGGTTTAGGTTTGGGAGGAGTTTCTTGACCTCTTGTTTATGAGTTTTTACGACTAAAATGCAATGCCAGTTTTCAGTCCGGGGACAAACTGCCTAGTTATGGATGAATTTTACCCTTTTTTTCCCCGTGTCTTTTTGGAATATTCTGCTTATTAATGCTTCCAATAACTAGCAGATGGAAAAAGGAAAAAGATAAAACTCAAATTCACTTTTTAATTCTTGTCTGTTATTATTTATTTGGGATCTCTCGTAATACTTTAAATTCTGAAAGTTTATTTCTGAAACCCAGTATTGTTATTCTCGGCTGCCCTGA |
| *Casp-3* target-2 | AAGGAATGACATCTCGGTCTGGTACAGATGTCGATGCAGCAAACCTCAGGGAAACATTCAGAAACTTGAAATATGAAGTCAGGAATAAAAATGATCTTACACGTGAAGAAATTGTGGAATTGATGCGTGATGGTAAGAAGAAACAATTAGAATGAACTTCATTGTACAATTAATTTTATACTATGGTCTAGCAATAATTATATGTATAATAAAATATGAGAATTGTGGAATTATACTGTACATAACCCTAATCTTACATACTTTAGTAAGAAGTTTAAAAAAAGTTATTTATGCCAACTTCCTAAAATGGTTTGAGATGTGTTGCCG |
| *EMX* target-1 | TATGGAAAAGAGCATGGGGCTGGCCCGTGGGGTGGTGTCCACTTTAGGCCCTGTGGGAGATCATGGGAACCCACGCAGTGGGTCATAGGCTCTCTCATTTACTACTCACATCCACTCTGTGAAGAAGCGATTATGATCTCTCCTCTAGAAACTCGTAGAGTCCCATGTCTGCCGGCTTCCAGAGCCTGCACTCCTCCACCTTGGCTTGGCTTTGCTGGGGCTAGAGGAGCTAGGATGCACAGCAGCTCTGTGACCCTTTGTTTGAGAGGAACAGGAAAACCACCCTTCTCTCTGGCCCACTGTGTCCTCTTCCTGCCCTGCCATCCCCTTCTGTGAATGTTAGACCCATGGGAGCAG |
| *EMX* target-2 | CCCTTCTGTGAATGTTAGACCCATGGGAGCAGCTGGTCAGAGGGGACCCCGGCCTGGGGCCCCTAACCCTATGTAGCCTCAGTCTTCCCATCAGGCTCTCAGCTCAGCCTGAGTGTTGAGGCCCCAGTGGCTGCTCTGGGGGCCTCCTGAGTTTCTCATCTGTGCCCCTCCCTCCCTGGCCCAGGTGAAGGTGTGGTTCCAGAACCGGAGGACAAAGTACAAACGGCAGAAGCTGGAGGAGGAAGGGCCTGAGTCCGAGCAGAAGAAGAAGGGCTCCCATCACATCAACCGGTGGCGCATTGCCACGAAGCAGGCCAATGGGGAGGACATCGATGTCACCTCCAATGACTAGGGTGGGCAACC |

**Table S10.** **On-target genotyping primers and Illumina adapter primers used for adding sequencing adapters to target amplicons**

| **Primer Name** | **Sequence (5’---3’)** |
| --- | --- |
| *CLTA* region-1_Miseq_F | GCTCCTCAGAGCACAGTTGTA |
| *CLTA* region-1_Miseq_R | GGATGCAGGGAGTCACAAATC |
| *CLTA* region-2_Miseq_F | TTTGGTAATCACTGTGTGCTATTT |
| *CLTA* region-2_Miseq_R | ATGGTTCGAGTGATGCGGAA |
| *HBB* target-1_Miseq_F | GCAATCATTCGTCTGTTTCCCA |
| *HBB* target-1_Miseq_R | CTGCTGGTGGTCTACCCTTG |
| *HBB* target-1_Miseq_F | GCAATCATTCGTCTGTTTCCCA |
| *HBB* target-1_Miseq_R | CTGCTGGTGGTCTACCCTTG |
| *HBB* target-3_Miseq F | CTGACCTCCCACATTCCCTTTT |
| *HBB* target-3_Miseq R | ATGGTTGGGATAAGGCTGGATT |
| *AIFM* target-1_Miseq_F | GGCTGACAACTTCTTATTGACC |
| *AIFM* target-1_Miseq_R | CCTTCCATAGAGAAGGCTGGAC |
| *AIFM* target-2_Miseq F | CGATCAGCTTAGCAGGTCACAG |
| *AIFM* target-2_Miseq R | TGTCCGTTTGGGAGTCACATCA |
| *Casp-3* taregt-1_Miseq_F | AGGCCTAGTAGGGTGTGTGA |
| *Casp-3* taregt-1_Miseq_R | TCAGGGCAGCCGAGAATAAC |
| *Casp-3* target-2_Miseq F | AAGGAATGACATCTCGGTCTGG |
| *Casp-3* target-2_Miseq R | CGGCAACACATCTCAAACCATT |
| *EMX* target-1_Miseq F | TATGGAAAAGAGCATGGGGCTG |
| *EMX* target-1_Miseq R | CTGCTCCCATGGGTCTAACATT |
| *EMX* target-2_Miseq F | CCCTTCTGTGAATGTTAGACCC |
| *EMX* target-2_Miseq R | GGTTGCCCACCCTAGTCATT |
| *CLTA* region-1_Adapter_F | TCGTCGGCAGCGTCAGATGTGTATAAGAGACAGGCTCCTCAGAGCACAGTTGTA |
| *CLTA* region-1_Adapter_R | GTCTCGTGGGCTCGGAGATGTGTATAAGAGACAGGGATGCAGGGAGTCACAAATC |
| *CLTA* region-2_Adapter_F | TCGTCGGCAGCGTCAGATGTGTATAAGAGACAGTTTGGTAATCACTGTGTGCTATTT |
| *CLTA* region-2_Adapter_R | GTCTCGTGGGCTCGGAGATGTGTATAAGAGACAGATGGTTCGAGTGATGCGGAA |
| *HBB* target-1_Adapter_F | TCGTCGGCAGCGTCAGATGTGTATAAGAGACAGGCAATCATTCGTCTGTTTCCCA |
| *HBB* target-1_Adapter_R | GTCTCGTGGGCTCGGAGATGTGTATAAGAGACAGCTGCTGGTGGTCTACCCTTG |
| *HBB* target-3_Adapter F | TCGTCGGCAGCGTCAGATGTGTATAAGAGACAGCTGACCTCCCACATTCCCTTTT |
| *HBB* target-3_Adapter R | GTCTCGTGGGCTCGGAGATGTGTATAAGAGACAGATGGTTGGGATAAGGCTGGATT |
| *AIFM* target-1_Adapter_F | TCGTCGGCAGCGTCAGATGTGTATAAGAGACAGGGCTGACAACTTCTTATTGACC |
| *AIFM* target-1_Adapter_R | GTCTCGTGGGCTCGGAGATGTGTATAAGAGACAGCCTTCCATAGAGAAGGCTGGAC |
| *AIFM* target-2_Adapter F | TCGTCGGCAGCGTCAGATGTGTATAAGAGACAGCGATCAGCTTAGCAGGTCACAG |
| *AIFM* target-2_Adapter R | GTCTCGTGGGCTCGGAGATGTGTATAAGAGACAGTGTCCGTTTGGGAGTCACATCA |
| *Casp-3* taregt-1_Adapter_F | TCGTCGGCAGCGTCAGATGTGTATAAGAGACAGAGGCCTAGTAGGGTGTGTGA |
| *Casp-3* taregt-1_Adapter_R | GTCTCGTGGGCTCGGAGATGTGTATAAGAGACAGTCAGGGCAGCCGAGAATAAC |
| *Casp-3* target-2_Adapter F | TCGTCGGCAGCGTCAGATGTGTATAAGAGACAGAAGGAATGACATCTCGGTCTGG |
| *Casp-3* target-2_Adapter R | GTCTCGTGGGCTCGGAGATGTGTATAAGAGACAGCGGCAACACATCTCAAACCATT |
| *EMX* target-1_Adapter F | TCGTCGGCAGCGTCAGATGTGTATAAGAGACAGTATGGAAAAGAGCATGGGGCTG |
| *EMX* target-1_Adapter R | GTCTCGTGGGCTCGGAGATGTGTATAAGAGACAGCTGCTCCCATGGGTCTAACATT |
| *EMX* target-2_Adapter F | TCGTCGGCAGCGTCAGATGTGTATAAGAGACAGCCCTTCTGTGAATGTTAGACCC |
| *EMX* target-2_Adapter R | GTCTCGTGGGCTCGGAGATGTGTATAAGAGACAGGGTTGCCCACCCTAGTCATT |

**Table S11. Predicted off-target sites and off-target amplicon sequences**

| **Off-Target Name** | **Chr** | **Off-Target Sequence (5’---3’)** | **Amplicon sequence (5’---3’)** |
| --- | --- | --- | --- |
| Cas9d-*HBB*-T3-OFFT1 | Chr6 | TTCC**a**TAAG**g**CCAACTACTA | TGCTTGCATGTAGTAGAACTTGTATTACGTCTATAGTAGGCTCATACATTCGAGAAAAAATAATTCAACCCCCAAAAGAT**GTGTAGTAGTTGGCCTTATGGAA**TATGCAAGATCATTAAATAAAAGTAGTAAGGATATCTGAATAAAGTCTAAATATCTATTTTATATCAATAAAGATATAATAAACCCACTTATTAAAGTTGGCTTAAATACTGTATTGAAAAAAAATTGTATGTTAAACACAAAAGATTCAACTTCAAAATAAAAAAAAAAATAGACAAGGAATCCCTAAGCAAATATTCACAAAAAGAAAGCTGAGTGGGTTTTCCACAGGCAGAGTGAGATCAATGAGATCCAGTGAAGCATG |
| Cas9d-*HBB*-T3-OFFT2 | chr10 | TT**t**CC**a**AAGTCC**c**ACTACTA | ATATGCCATAGGACATGGATCCAAATATTTACTTTTCTCTTAGTTACACATACTTTTATCTATATCTTTTATCTTACCTTTACTTTATGGCCAATTTATTGATTTTATAGTAATAATTTATGTTTTTAGGCTTTAAAGGAAGTATATAAACATTTAATTATATATTTGCTTAACAGTACAGCCTATTTTATTAGGCAATTTAAAATTTTTAACTTTCTGCTGAACA**TTTCCAAAGTCCCACTACTATAC**AAATTAATTAGTTAAGAAATGGAATTCATTGTCGTTATTATTTTAAATATATTCTCTCTCCATCCCTATCTTAGTTAATGATACTATCACTCTTTTAGGCTCAATTTCAAAAGCATCTTCTTATTCCATCTTTTCCCTCTGCCAGTATATTCAATTCCA |
| Cas9d-*AIFM*-T1-OFFT1 | Chr17 | TGTGTG**tg**TGTACCAGG**g**AG | TGAGCAGTTTCGACAGGGGTTTATTTAAATTCTTATTTATTTTTAAAAATGAAAATGTAAATGTGATTTGCTATTAGCTCTGCTCTTAAACAGCACGGGACTCACCTGTCAGATTTCCTGGAGTGAGGGGGATGATGGTGCAGTGCCAGTCGTAGAATGGGGTGGGCCTGTTTCTGATCTAGAGAGGTCTACGGGGTTCCCCTGCACACCAGGCAACCACAGGGCCTGCAGAACTGCTTG**TGTGTGTGTGTACCAGGGAGCAC**ATAGGTGGCACTCCCTACGTGGTGCCCTCAGACCGTGATGTTGCAAAGCA |
| Cas9d-*AIFM*-T1-OFFT2 | chr1 | TGTGTGA**t**TGTACCAG**a**T**t**G | CATGGGCAATTGCAGTCCAGATGGGATGCAGAGAGGGGGCCTGGCATTAGACCAGCACCATCCTACTGCACAACCAGTTGGCACTGGCTGCACTCCTGTTTTTGCTCAAGATAGGAAATAGTTTCTGGGTCCGGGCCTGGGCGAACAGTCCTCTGGGCCAGAGCCATAGAGAACTATACCTGTGTC**TGTGTGATTGTACCAGATTGGAC**TCTCTTTCAATGTATATTTTGGTCTGTGTTTACAGAGAATATCGTGTCCAAGATGGAAAAATCTTCATTCCCTTGAACATCACTGTGCAAGGAGACGTG |
| Cas9d-*AIFM*-T1-OFFT3 | Chr16 | TGTGTG**g**CTGTACCA**at**TAG | TCCGAGGCTTCGTTCCAATCACCAAGGCTACCAAGGCCACTGGGAGGGACAAACAGGGAGAGGCCTGCACCTGCTTCCAAAAGGACCTCCGTGCCAGGGTAGGCGAACGGTGGAAGCACAGTG**TGTGTGGCTGTACCAATTAGAAC**ATGCAATTACCCAGGCTGCCAATGCAAATAGACCTAACAGAGGCCCCTCGTCCAGCTCATCTGCATGCACAAAAGGTGGCTGGCCAGCACTTAGAAACCCGTGGACCACGCTGGACGCAGGCAGGCCCTGCTTTAACAGAACCCTAAGTAAAAAGCCCTTTGCAATGTAGCATCATGTTAATGGGATGCAAAACAGCACAGGGTGAGGCAGGAAGGAGTGAGACTGATAGGGCTGGGTTGGAA |
| Cas9d-*AIFM*-T2-OFFT1 | Chr8 | CA**tt**TTAGATAACACCTA**t**T | CCTCACAGGACTCTGATCCTAGCTCTCTGTTCGCTAATTACTACTGGTATCTGTTAACTCATGATCCTTTTCAGCAGCATTTTCAAAATCCTAAACTGCTCCAAGGGAAATGAAAAGGCTTCATGGCAAAACAAGTTTGAAAACTTTGAAAATCCAGGTCTCACTCTGAGAATCAAAATAGATACCTACACAGATGCATAAAGCCTCTGAACATTCCACAGTAAAGTGACTTGTGTAAACATTGTTTCACCCAGCATTGCTCAAACATATTTAATTATGGAAATT**CATTTTAGATAACACCTATTAAC**ATCTTATGGATTAGTTTTCTCCATATTCTAACTCACTTCAATTGAGCAAACACATTTACAGAAACTACTTGTTGCATCAAGGAACTCATAAGTGAGAGCAGTGAT |
| Cas9d-*AIFM*-T2-OFFT2 | Chr3 | CAACTTA**t**ATAACA**a**C**c**AGT | AACTGGCTTCAGTGTAAATACTAGAAGCCACAAAATAACATATTGAAAATTTAACTACGTAAGAATAAAACTGTACACACACACACACACACACACACACACACACACACACACGGAATGTCCAAAGCCAAATAACAAAGGGAAACAATTTG**CAACTTATATAACAACCAGTAAC**CACTTTTCTTAATACATAAAAGAACTCATAAGAATCAGAAAGAAAATGGCCAAGAACACAACAGAATTGATCAAAAGACACAAACCATTTTCAGATAAAGAAATGCAAATGACTATAAGGTATGTGAAGATACACAAACTCATTAATTATAAGAAAATACAAGTTAAAATTATAGGGTGATAGTTCATTTTTACTCTCAAATAGGGAATAATAAAAAATTTGATCCATACATTTCGCTGATGT |
| Cas9d-*AIFM*-T2-OFFT3 | Chr4 | **gc**AC**a**TAGA**a**AACACCTAGT | TGGTGAGGTTAATGACAGAGGCTACTCTTCAGATTTCAAATTTCAGCTTACTGATATTTAGTTTTCAGACAAATAAATATTAACATTGGATGTTTTCCTGTATAGCAGTGGTAAAATTGTAAATTTTATTTAATGTTAATACAGGTAGCTAGT**GTTACTAGGTGTTTTCTATGTGC**TAGGCATTGTTCTAAGTATTTAACATGTCTTATTTTCTTTAAGCCTCTCAACAATCTTTTCAGCTGGTATTATTACCAAATCTCTTTTAGATATGATAAATCAGAGGCAGTATAATGACCTTAAGTTTACCTCTCCAAGTTACACACTTGGGCACTAAGGGC |
| Cas9d-*Casp3*-T1-OFFT1 | Chr6 | AGAA**t**C**c**CaTCCCAAACCTA | TGTGATGAAGGACTTTTCAGGCAGAGGTAACAATATTAACACAGTGCCTGGGTACTTAGTAAAAACGGAGTAACTGATGAGAGTGTTGGTTATTGTTGTGTTGGTGGTGATGATACAGTTTTAGGGG**GTATAGGTTTGGGATGGGATTCT**TAGAAAGATGGTAGATTTAAGCGTATTCATAGCTTGTGGGGACATAACTAGTAGAGTGTAAAAATACAGCATAGAAGGAATGAAGGACTGGAAAGAAAGAGAAACTCCAGAGTACAACCAATGTGGGTTA |
| Cas9d-*Casp3*-T1-OFFT2 | chr17 | **c**GAAA**a**TCCTCCCAA**g**CCTA | AGTCGGCCAGGAGAGAAAGAAACAGTCATGCCAGAACCTGGAGGCCCCTCCCTGACAGGTGCAGACCATGTTGAGCATGGAGCTGTGGCTTTTATGTTGAGCACGGAGGGGTGGGTTTATCGCCACCCTCGCTGACAATGTCACTATTCCTCTGTTCTGCCGCCGCCAGCCAATGGGCATGGCCAG**GTGTAGGCTTGGGAGGATTTTCG**ATGCCTGGACTCGACTCTCTGCCTCCAGTTGGGGAGGACAAAGAGCGCAGCTTGTCCCGGCCCTGGGACCGTCCTACCAGGGAAACACAGGGCAGAGGCCATGAAGTAGACATGAAATGGCCGGGTTCA |
| Cas9d-*Casp3*-T1-OFFT3 | chr19 | AGAAAC**c**CCTCCCA**g**ACCT**g** | CCCTGGTCCTCATGGACAATGGCAGGTGCTGATGGCTGTGGCTGTGTGACAGCCATGGAGGTGGGGTGGGTTCTGGACCTCAGTGGCCAGGGCAGAAGCAGGACATGCTATGTGCATCCAGAACCAATTGAGATGATGTCTCCAACACGACAGAACAGGAGGTGGCTCGCAGGTTCA**GTCCAGGTCTGGGAGGGGTTTCT**CATCTCTGGGTGTCACTCCTGCTTCCTTTTTCCTCCTCTAGTTCCTCAACACTGTTGTTACAAAGTCCCTGCATTTAACTCCCTCCATCTGAAACCCTCCAAGTGGTTTCCGTCTTCT |

**Bold pink: mismatched nucleotides**

**Bold blue: off-target sequences**

**Bold blue: PAM sequences**

**Table S12. Off-target PCR primer sequences**

| **Primer Name** | **Sequence (5’---3’)** |
| --- | --- |
| Cas9d-*HBB*-T3-OFFT1-F | TGCTTGCATGTAGTAGAACTTGTA |
| Cas9d-*HBB*-T3-OFFT1-R | CATGCTTCACTGGATCTCATTGATC |
| Cas9d-*HBB*-T3-OFFT2-F | ATATGCCATAGGACATGGATCCAA |
| Cas9d-*HBB*-T3-OFFT2-R | TGGAATTGAATATACTGGCAGAGGG |
| Cas9d-*AIFM*-T1-OFFT1-F | TGAGCAGTTTCGACAGGGGT |
| Cas9d-*AIFM*-T1-OFFT1-R | TGCTTTGCAACATCACGGTCTGAG |
| Cas9d-*AIFM*-T1-OFFT2-F | CATGGGCAATTGCAGTCCAGATG |
| Cas9d-*AIFM*-T1-OFFT2-R | CACGTCTCCTTGCACAGTGATG |
| Cas9d-*AIFM*-T1-OFFT3-F | TCCGAGGCTTCGTTCCAATCAC |
| Cas9d-*AIFM*-T1-OFFT3-R | TTCCAACCCAGCCCTATCAGTCT |
| Cas9d-*AIFM*-T2-OFFT1-F | CCTCACAGGACTCTGATCCTAGC |
| Cas9d-*AIFM*-T2-OFFT1-R | ATCACTGCTCTCACTTATGAGTTCC |
| Cas9d-*AIFM*-T2-OFFT2-F | ACTGGCTTCAGTGTAAATACTAGAAG |
| Cas9d-*AIFM*-T2-OFFT2-R | ACATCAGCGAAATGTATGGATCAA |
| Cas9d-*AIFM*-T2-OFFT3-F | TGGTGAGGTTAATGACAGAGGCTAC |
| Cas9d-*AIFM*-T2-OFFT3-R | GCCCTTAGTGCCCAAGTGTGTA |
| Cas9d-*AIFM*-T1-OFFT1-F | TGAGCAGTTTCGACAGGGGT |
| Cas9d-*Casp3*-T1-OFFT1-F | TGTGATGAAGGACTTTTCAGGCAG |
| Cas9d-*Casp3*-T1-OFFT1-R | TAACCCACATTGGTTGTACTCTGG |
| Cas9d-*Casp3*-T1-OFFT2-F | AGTCGGCCAGGAGAGAAAGAA |
| Cas9d-*Casp3*-T1-OFFT2-R | TGAACCCGGCCATTTCATGTC |
| Cas9d-*Casp3*-T1-OFFT3-F | CCCTGGTCCTCATGGACAATGG |
| Cas9d-*Casp3*-T1-OFFT3-R | AGAAGACGGAAACCACTTGGAGG |
| Illumina-Cas9d-*HBB*-T3-OFFT1-F | TCGTCGGCAGCGTCAGATGTGTATAAGAGACAGTGCTTGCATGTAGTAGAACTTGTA |
| Illumina-Cas9d-*HBB*-T3-OFFT1-R | GTCTCGTGGGCTCGGAGATGTGTATAAGAGACAGCATGCTTCACTGGATCTCATTGATC |
| Illumina-Cas9d-*HBB*-T3-OFFT2-F | TCGTCGGCAGCGTCAGATGTGTATAAGAGACAGATATGCCATAGGACATGGATCCAA |
| Illumina-Cas9d-*HBB*-T3-OFFT2-R | GTCTCGTGGGCTCGGAGATGTGTATAAGAGACAGTGGAATTGAATATACTGGCAGAGGG |
| Illumina-Cas9d-*AIFM*-T1-OFFT1-F | TCGTCGGCAGCGTCAGATGTGTATAAGAGACAGTGAGCAGTTTCGACAGGGGT |
| Illumina-Cas9d-*AIFM*-T1-OFFT1-R | GTCTCGTGGGCTCGGAGATGTGTATAAGAGACAGTGCTTTGCAACATCACGGTCTGAG |
| Illumina-Cas9d-*AIFM*-T1-OFFT2-F | TCGTCGGCAGCGTCAGATGTGTATAAGAGACAGCATGGGCAATTGCAGTCCAGATG |
| Illumina-Cas9d-*AIFM*-T1-OFFT2-R | GTCTCGTGGGCTCGGAGATGTGTATAAGAGACAGCACGTCTCCTTGCACAGTGATG |
| Illumina-Cas9d-*AIFM*-T1-OFFT3-F | TCGTCGGCAGCGTCAGATGTGTATAAGAGACAGTCCGAGGCTTCGTTCCAATCAC |
| Illumina-Cas9d-*AIFM*-T1-OFFT3-R | GTCTCGTGGGCTCGGAGATGTGTATAAGAGACAGTTCCAACCCAGCCCTATCAGTCT |
| Illumina-Cas9d-*AIFM*-T2-OFFT1-F | TCGTCGGCAGCGTCAGATGTGTATAAGAGACAGCCTCACAGGACTCTGATCCTAGC |
| Illumina-Cas9d-*AIFM*-T2-OFFT1-R | GTCTCGTGGGCTCGGAGATGTGTATAAGAGACAGATCACTGCTCTCACTTATGAGTTCC |
| Illumina-Cas9d-*AIFM*-T2-OFFT2-F | TCGTCGGCAGCGTCAGATGTGTATAAGAGACAGACTGGCTTCAGTGTAAATACTAGAAG |
| Illumina-Cas9d-*AIFM*-T2-OFFT2-R | GTCTCGTGGGCTCGGAGATGTGTATAAGAGACAGACATCAGCGAAATGTATGGATCAA |
| Illumina-Cas9d-*AIFM*-T2-OFFT3-F | TCGTCGGCAGCGTCAGATGTGTATAAGAGACAGTGGTGAGGTTAATGACAGAGGCTAC |
| Illumina-Cas9d-*AIFM*-T2-OFFT3-R | GTCTCGTGGGCTCGGAGATGTGTATAAGAGACAGGCCCTTAGTGCCCAAGTGTGTA |
| Illumina-Cas9d-*Casp3*-T1-OFFT1-F | TCGTCGGCAGCGTCAGATGTGTATAAGAGACAGTGTGATGAAGGACTTTTCAGGCAG |
| Illumina-Cas9d-*Casp3*-T1-OFFT1-R | GTCTCGTGGGCTCGGAGATGTGTATAAGAGACAGTAACCCACATTGGTTGTACTCTGG |
| Illumina-Cas9d-*Casp3*-T1-OFFT2-F | TCGTCGGCAGCGTCAGATGTGTATAAGAGACAGAGTCGGCCAGGAGAGAAAGAA |
| Illumina-Cas9d-*Casp3*-T1-OFFT2-R | GTCTCGTGGGCTCGGAGATGTGTATAAGAGACAGTGAACCCGGCCATTTCATGTC |
| Illumina-Cas9d-*Casp3*-T1-OFFT3-F | TCGTCGGCAGCGTCAGATGTGTATAAGAGACAGCCCTGGTCCTCATGGACAATGG |
| Illumina-Cas9d-*Casp3*-T1-OFFT3-R | GTCTCGTGGGCTCGGAGATGTGTATAAGAGACAGAGAAGACGGAAACCACTTGGAGG |
